# Layer 6 ensembles can selectively regulate the behavioral impact and layer-specific representation of sensory deviants

**DOI:** 10.1101/657114

**Authors:** Jakob Voigts, Christopher A Deister, Christopher I Moore

## Abstract

Predictive models can enhance the salience of unanticipated input, and the neocortical laminar architecture is believed to be central to this computation. Here, we examined the role of a key potential node in model formation, layer (L) 6, using behavioral, electrophysiological and imaging methods in mouse somatosensory cortex. To test the contribution of L6, we applied weak optogenetic drive that changed which L6 neurons were sensory-responsive, without affecting overall firing rates in L6 or L2/3. This stimulation suppressed L2/3 deviance encoding, but maintained other stimulus encoding. The stimulation also selectively suppressed behavioral sensitivity to deviant stimuli without impacting baseline performance. In contrast, stronger L6 drive inhibited firing and suppressed overall sensory function. These findings indicate that, despite their sparse activity, specific ensembles of stimulus-driven L6 neurons are required to form neocortical predictions, and for their behavioral benefit.

## Introduction

The six-layered architecture of mammalian neocortex emerged relatively late in evolution. A common proposal is that this structure supports formation of complex predictive models, thereby enabling the rapid and adaptive shifts in behavior underlying capabilities such as flexible language. An elemental example of model formation is evident in the response to deviations from ongoing patterns. When identical sensory stimuli are repeated and then a ‘deviant’ occurs, neocortical neurons often fire differently than they would to the deviant in isolation, or after its repetition^1,2^. Signatures of this computation are found in visual^3^, auditory^4,5^, and language processing^6^ areas. Such change detection is typically studied as the increase in firing rates elicited by deviant stimuli following stimulus-specific adaptation^7^. Stimulus-tuned neurons along the afferent pathway, including at thalamocortical synapses, adapt to repeated stimulation^8–10^ and subsequent deviant stimuli activate new pools of less adapted neurons, leading to increased neocortical drive.

However, in addition to such bottom-up adaptation mechanisms, neocortically-represented factors such as stimulus context, history, and expectation^11–14^ also influence sensory responses, supporting the hypothesis that neocortical representations could be key to deviant processing. While likely all neocortical layers contribute to model implementation, layers 2/3 (L2/3) are a leading candidate for the representation of more complex information. These supragranular layers are typically the first to express receptive field plasticity in response to sensory change, prior to L4^15^, and are more susceptible to modulation by shifts in attentional state than deeper layers^16^. Further, in primary somatosensory and visual neocortices, L2/3 receptive fields can encode specific temporal stimulus patterns^17,18^, and can integrate multiple types of information (e.g., motor and sensory signals) and the mismatch in their alignment^19^.

Layer 6 (L6) is also well positioned to contribute to the neocortical implementation of predictive models, as it integrates lemniscal thalamic, long-range cortico-cortical, and modulatory inputs^20–24^. Corticothalamic L6 neurons (CT) in primary visual and somatosensory neocortex are sparsely sensory driven^25^ with selective receptive fields^24^, and can robustly modulate sensory gain through an intracortical pathway^26^. These findings suggest that specific populations of L6 neurons could regulate sensory responses depending on stimulus context. This prediction is supported by L6-mediated modulation of visual receptive fields by stimulus context^27^ and by preferential involvement of deep cortical layers in top-down sensory processing^28^. However, whether L6 contributes to the representation of stimulus changes across layers, and to perception, is not known. Here, we tested this hypothesis using selective optogenetic modulation of L6 activity in awake behaving mice, single-neuron recordings across neocortical layers, and 2-photon Ca^2+^ imaging.

## Results

### Weak drive of L6 CT impaired the behavioral detection of deviant, but not baseline stimuli

We first tested the impact of manipulating the activity of L6 CT cells on change detection behavior in a naturalistic and untrained sensory decision-making task, gap-crossing^29^ (Fig. 1 a). In this task, mice use their vibrissae to locate and cross between elevated platforms whose distance is changed after each trial (~4-6 cm, 6 mice, Extended Data Figure 1). Experiments were performed under near infrared illumination and auditory white noise to reduce visual or auditory confounds^29,30^. Only trials with crossings within 5 seconds of exploring the gap were analyzed. We applied selective optogenetic depolarization to L6 in mice expressing Channelrhodopsin (ChR2) in L6 corticothalamic (CT) pyramidal cells (GN 220-NTSR1 Cre line^31^).

**Figure 1.**
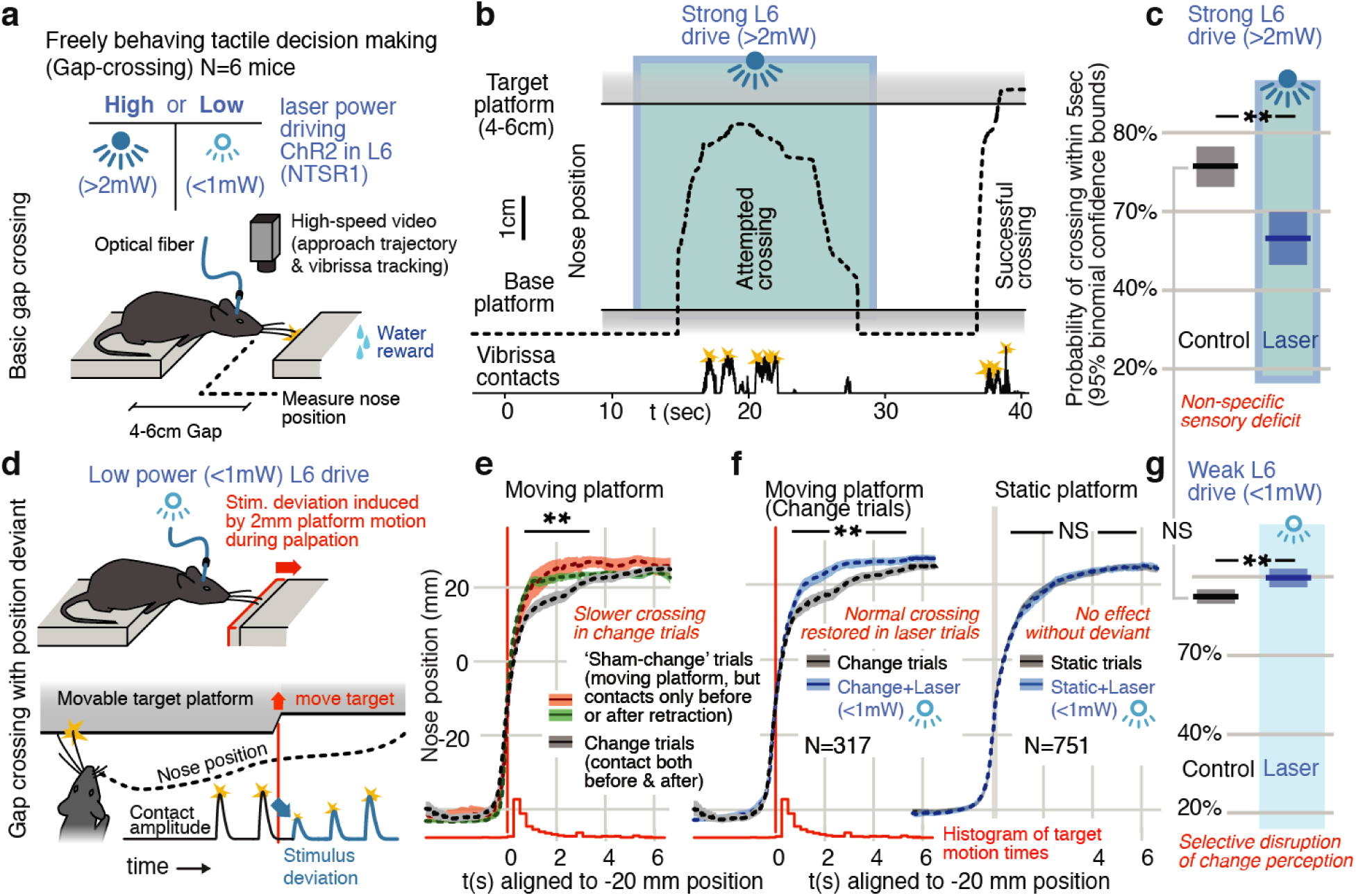
In a natural behavior, strong optogenetic drive of L6 causes a sensory deficit, while weak drive selectively negates detection of deviant stimuli. **(a)** To test the impact of L6 modulation in a naturalistic context, we used an unrestrained gap-crossing task and applied high or low power optogenetic drive to L6 CT cells (ChR2 in NTSR-1 cre line). **(b)** Example trace (nose position over the gap) where strong optogenetic L6 drive disrupted gap-crossing behavior. **(c)** Strong L6 drive made mice less likely to cross the gap (N=514 trials, bootstrapped 95% confidence intervals (CIs)). **(d)** To create a sudden, small sensory deviation, the target platform was retracted by ~2 mm during bouts of tactile sampling^32^. **(e)** When the platform was retracted in the middle of a bout of vibrissal contacts, mice slowed their approach relative to trials in which they contacted the target either only before (*red*) or only after (*green*) platform retraction. **(f)** Weak L6 drive removed the deceleration associated with platform retraction (*left*), but had no effect when the platform was static (*right*), showing that it selectively abolished change detection with little effect on other sensory and sensorimotor function. See Extended Data Figure 3 for per-mouse data. **(g)** Weak L6 drive also did not reduce crossing probability (N=751 trials, analysis as in c).

We tested mice on the gap-crossing task under two conditions: strong L6 CT optogenetic activation (>2mW power) which recruits trans-laminar inhibition and reduces neocortical sensory gain in V1^26^, or weak L6 drive (<1mW) (Fig.1 a). The strong drive led to non-specific sensory deficits, reducing the likelihood of successful gap-crossing from ~75% to ~50% (P<0.01, Fig. 1b,c), but had no detectable effect on the whisking pattern of the mice (Extended Data Figure 2).

**Figure 2.**
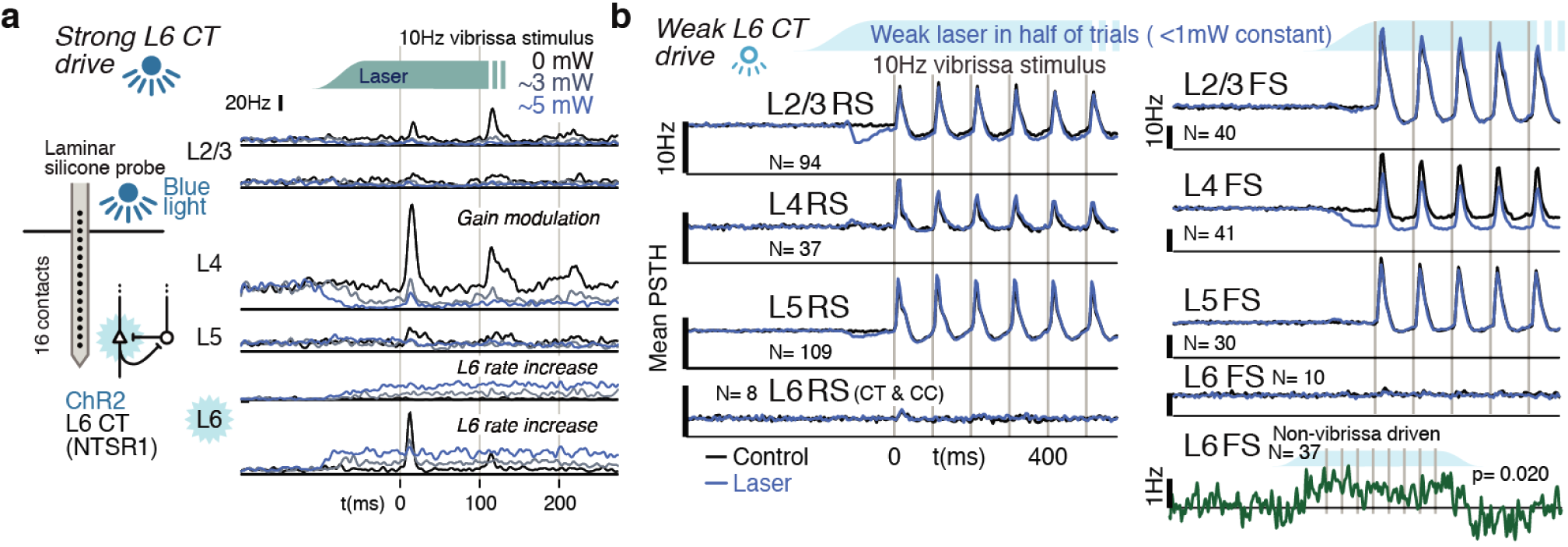
Weak optogenetic stimulation of L6 CT does not significantly impact overall regular firing rates in RS cells across layers. **(a)** Strong optogenetic L6 CT drive increased L6 firing rates and reduced sensory gain in other layers, as in visual cortex^26,38^. Circuit diagram adapted from^26,40^. **(b)** Low power (<1mW) optogenetic drive did not change mean sensory evoked firing rate of RS in any layer, and small FS firing rate changes were observed in only L6 FS that were not sensory responsive (increased activity) and sensory-responsive L4 FS (decreased activity).

To test the effect of L6 drive on change detection, we induced small stimulus deviations during gap-crossing: In ~35% of trials the target platform was rapidly pulled back by ~2mm during a bout of exploration^32^ (Fig.1d). To control for non-specific cues that could have been associated with retraction (e.g., sound or air currents), true ‘change’ trials in which the vibrissae contacted the retracting platform both before and after the retraction (as illustrated in Fig.1 d, N = 317) were compared to ‘sham-change’ trials in which they contacted the platform either only before (N = 49) or only after (N = 134) a platform motion. On these trials, mice could not perceive a change in position through vibrissal palpation, but would experience any other effects of platform motion. In the true change condition, but in neither of the sham conditions, mice slowed their approach (Fig. 1e, P<0.005 change vs. only before, and P<0.01 change vs. only after, rank sum), presumably to precisely re-locate the target before crossing, showing that mice perceive and react to the sensory deviant.

We next tested whether weaker L6 optogenetic drive (<1mW) would impair this change detection behavior. Indeed, weak L6 drive removed the extra sampling time that platform motion would otherwise generate (P<0.001 laser vs. control, N=317 trials, Fig. 2f left, Supplementary Movie 1). Weak L6 drive did not affect behavior when the target platform was static (P=0.9, N=751 trials, Fig. 2f right, Extended Data Figure 3). The weaker drive also slightly increased the overall likelihood of crossing (Fig.1 g, same analysis as Fig.1 c). Gap-crossing is sensitive to changes in sensory function^33,34^, the maintenance of regular gap-crossing behavior in the static platform condition therefore indicates that the effects of weak L6 CT drive are specific to change detection. In separate experiments using head-fixed vibrissa stimulus detection with upward vs. downward deflection deviants (Extended data figure 4), we found similar disruption of change detection, and not of overall sensitivity with weak L6 drive. Disrupting L6 activity through weak optogenetic drive therefore selectively disrupts behavioral detection of stimulus changes.

**Figure 3.**
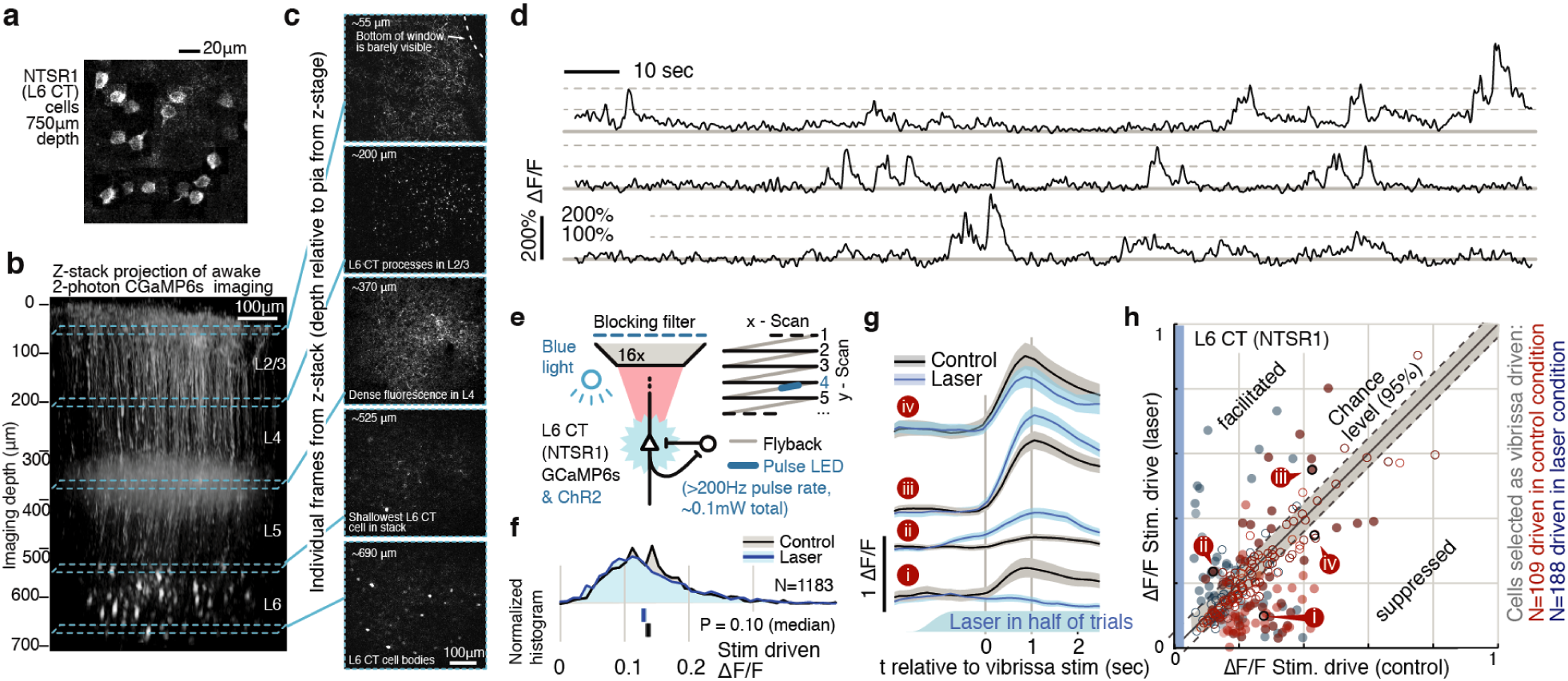
Weak depolarization of L6 CT maintains overall rates but changes the identity of stimulus-driven ensembles. To test the impact of weak optogenetic drive on L6 CT, we combined optogenetic stimulation with awake 2-photon imaging using GCaMP6s expression in the L6 specific NTSR1-Cre line (see Methods). **(a)** Example image L6 CT cells, shown as sum over 12 individual frames, in each frame a subset of the cells were active. **(b)** Z-stack projection from GCaMP6s expression in L6 CT. **(c)** Sample frames from different depths of the Z-stack. **(d)** Example time series showing typical signal-to-noise ratios. **(e)** Optogenetic activation was interleaved with imaging at >200Hz during laser scanning ‘flyback’. **(f)** Optogenetic drive did not significantly change the mean population response of L6 to sensory input, ranksum test, CI via bootstrap of median. **(g)** However, individual L6 cells showed facilitation or suppression of vibrissa-evoked responses during optogenetic drive. **(h)** Optogenetic L6 activation did not change the overall output from L6, but shifted the identity of the activated ensemble (P<0.001 IQR vs. shuffled control), regardless of whether cells were classified as vibrissa-driven in the control (*red*) or in the laser (*blue*) condition (Extended Data Figs. 9, 10). Filled circles indicate cells where the laser effect was individually significant per-cell (P<0.0025, at 1 sec time point, via bootstrap). 25.0% of cells that were vibrissa driven in the laser condition were significantly facilitated, 23.9% suppressed. For cells driven in the control condition, 10.9% were facilitated, 32.9% suppressed.

**Figure 4.**
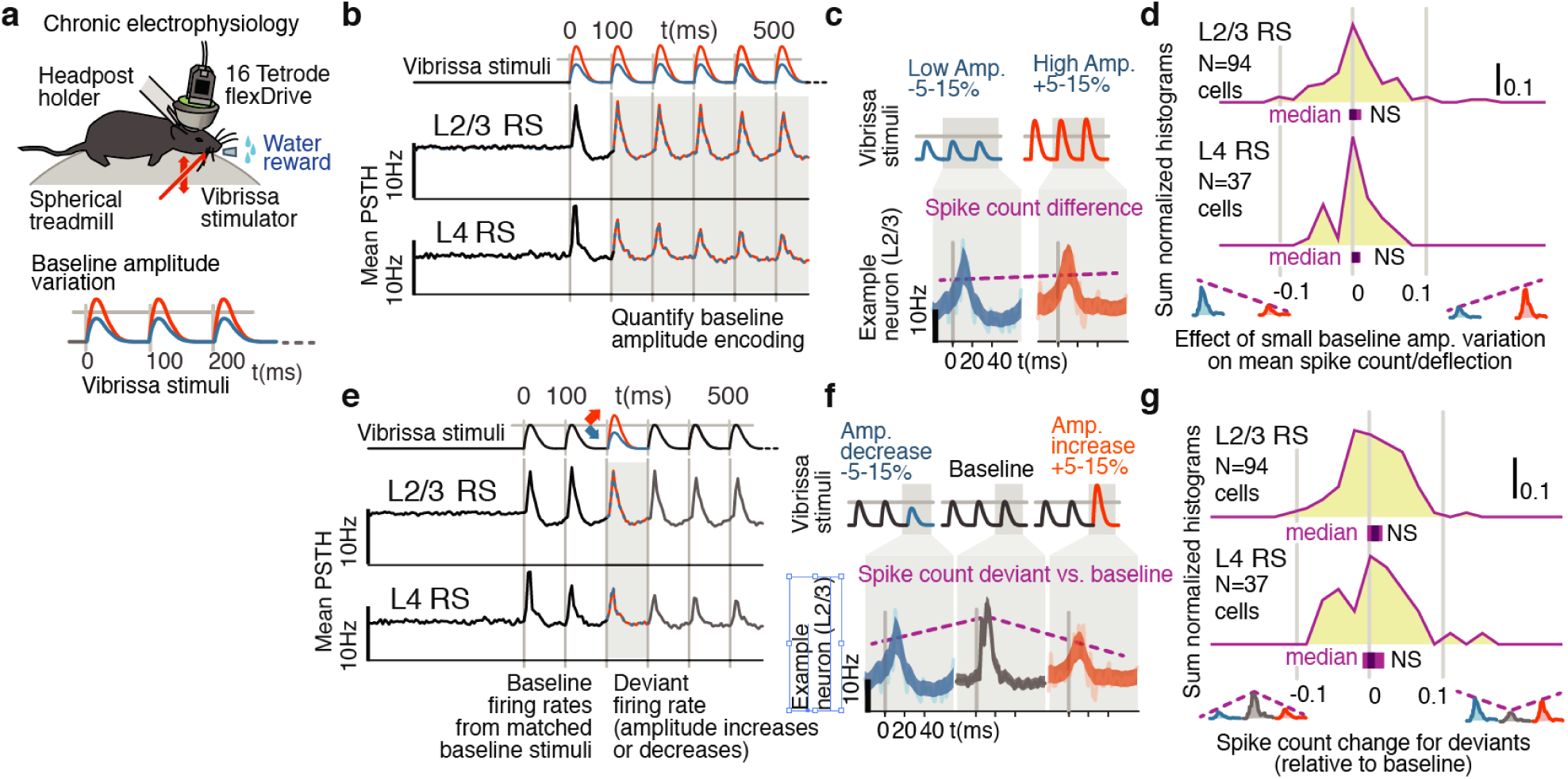
Small variations in stimulus amplitude were not robustly encoded in layers 4 and 2/3 SI. **(a)** Experimental preparation. Baseline stimulus amplitudes were varied by up to 15% on a trial-by-trial basis, in a range of ± ~25mm/second, and presented in trains of 7 stimuli at 10 Hz. **(b)** The average sensory-evoked PSTHs from sensory-responsive RS neurons show that neither L2/3 nor L4 cells significantly changed their mean firing rates to reflect stimulus amplitude in the range employed here (95% confidence intervals (CI) are shown by the width of the traces). **(c)** Changes in spike rate in an example neuron. **(d)** The population distribution of differences in spike rates per deflection between small and large baseline stimulus amplitudes showed no significant encoding of amplitude. *Purple bars* beneath each histogram indicate 95% Cis (via bootstrap of median), ranksum test. **(e)** To probe the encoding of changes, rather than of overall amplitude, we presented amplitude deviations in the middle of stimulus trains. **(f)** Example of a neuron where stimulus amplitude deviants of either increasing or decreasing amplitude caused decreases in firing. **(g)** To test for generalized sensitivity to deviants, we calculated the net increase or decrease in firing to both types of amplitude deviants relative to the baseline response. While a few individual cells showed a generalized sensitivity to deviation, amplitude deviants did not systematically affect overall firing rates in L2/3 or L4, as shown in the centered population distribution.

### Weak depolarization of L6 neurons changed the identity of stimulus-driven ensembles in L6 without changing mean firing rates across layers

We next tested the impact of strong (disrupts gap-crossing) and weak (selectively disrupts change detection) optogenetic L6 drive regimes on sensory encoding. To approximate the stimulus statistics that occur during gap-crossing in head-fixed mice, we delivered trains of vibrissa deflections (7 deflections, 10 Hz) to the B and C row vibrissae, arcs 1-3, and recorded in matched somatotopic positions in vibrissal primary somatosensory neocortex (SI). Strong L6 drive reduces sensory gain^26,35^ in V1 and we observed analogous suppression of sensory responses in SI, when we drove L6 rates above their baseline (Fig.2a, acute laminar probe recording).

We next investigated sensory responses under weak L6 drive using chronic tetrode recordings in awake mice. To characterize layer-specific activity, we implanted high-density arrays of movable multi-contact electrodes^36^ in SI, yielding ~25 identified neurons per session (5 mice, Extended Data Figure 5). We recorded 1242 neurons over 75 sessions, 395 of which were phasically stimulus-driven (see Methods). To classify neurons by layer, we tracked electrode depth and stimulus-evoked LFP^37^, and categorized their spike waveform^38^ as regular spiking (‘RS’; typically excitatory pyramidal neurons) or fast-spiking (‘FS’; typically inhibitory interneurons: ~30% were FS). Consistent with prior studies^22,25,39^, L6 neurons were sparsely sensory responsive, as only 18 (RS and FS) of 139 (~13%) were phasically sensory driven. Further, driven L6 neurons were also sparse in their response rates, showing only small vibrissa-driven increases in their activity (Fig. 2b).

**Figure 5.**
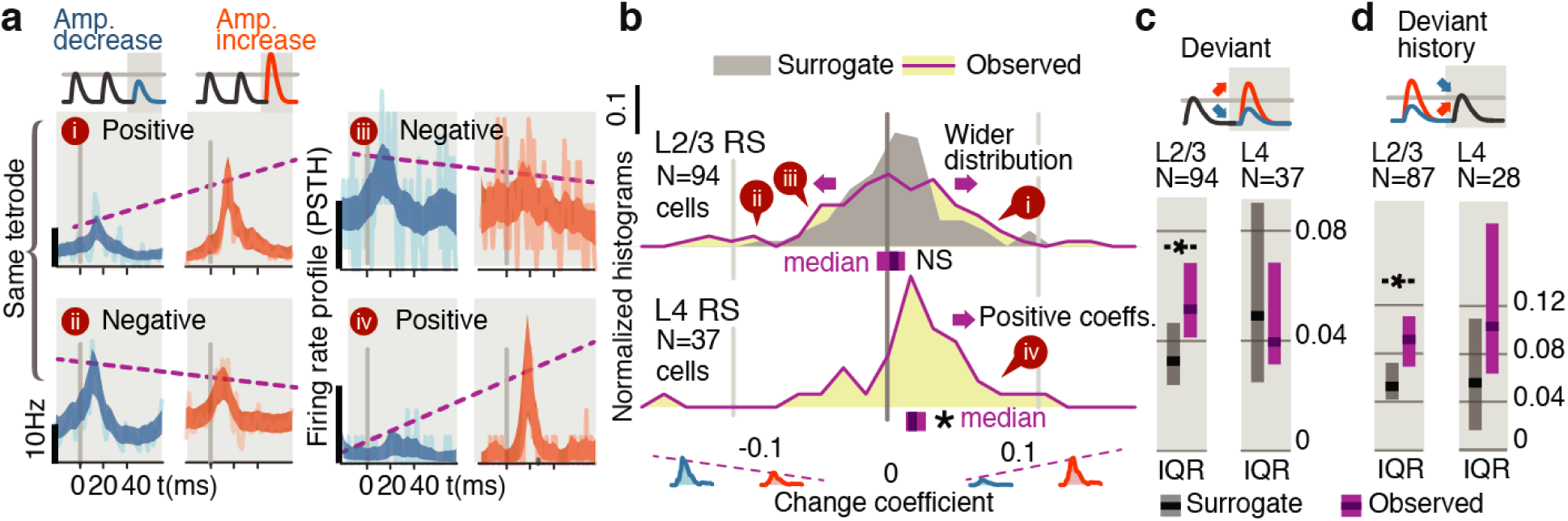
Small stimulus deviations were represented by distinct patterns of rate changes in layers 4 and 2/3. **(a)** Stimulus amplitude deviants were presented after 2-6 repeated baseline amplitude stimulations. Examples show sensory-driven PSTHs, shaded regions indicate the 95% CI. *Red traces* are responses to amplitude increases, and *blue* to amplitude decreases. We calculated a ‘change coefficient’, defined as spike count differences between responses to increased versus decreased amplitudes (*purple dashed line*). Neurons had diverse responses, including positive change coefficients (higher firing probability for stimulus increases - i, iv) and negative change coefficients (higher probability for stimulus decreases - ii, iii). **(b)** Histograms of the distribution of change coefficients for all stimulus-driven RS in L4 and L2/3. The L4 population responded to stimulus changes with positive change coefficients. In contrast, L2/3 showed several types of responses to deviants, with both positive and negative change coefficients, reflected in a broadening of the change coefficient distribution (*purple*) relative to a surrogate distribution of shuffled baseline stimuli (*gray*). Bars show the 95% CIs (*purple*) and the median value. **(c)** The L2/3 population, but not L4, was broader than a surrogate distribution, quantified via the interquartile range (IQR), ranksum test, bars graphs show median and 95% CI via bootstrap of median. **(d)** Heterogeneous encoding (reflected in broadening of the distribution) was also observed in L2/3 when initial baseline amplitudes were larger or smaller than a deviant that always had the same intermediate amplitude, demonstrating encoding of deviants from stimulus history and not absolute amplitude encoding in L2/3.

We applied weak L6 drive as during gap-crossing (~0.1-0.5 mW total), and ramped the light intensity over >100 msec prior to sensory stimulation, to prevent overlap of sensory responses and any onset transients. In contrast with the stronger drive, this stimulus did not impact sensory evoked firing rates in RS from any layer (Fig. 2b, Extended Data Figure 6). The only significant impact on sensory responses was a modest decrease in sensory responses in L4 FS (31.7 vs. 24.6 Hz median peak rates, P=0.001, Fig. 2b). A small increase in firing rates in non-sensory responsive L6 FS was also observed (Fig. 2b, P=0.02 sign-rank, laser - control firing rates, N=37). Increased activity in these L6 FS presumably offset the direct effects of optogenetic drive in L6 RS, keeping L6 RS firing rates at a baseline level (P=0.18, N=89, signed rank). In L2/3 and L5 RS, transient suppression was evident after optogenetic stimulus onset, but rates returned to baseline prior to sensory stimulation.

**Figure 6.**
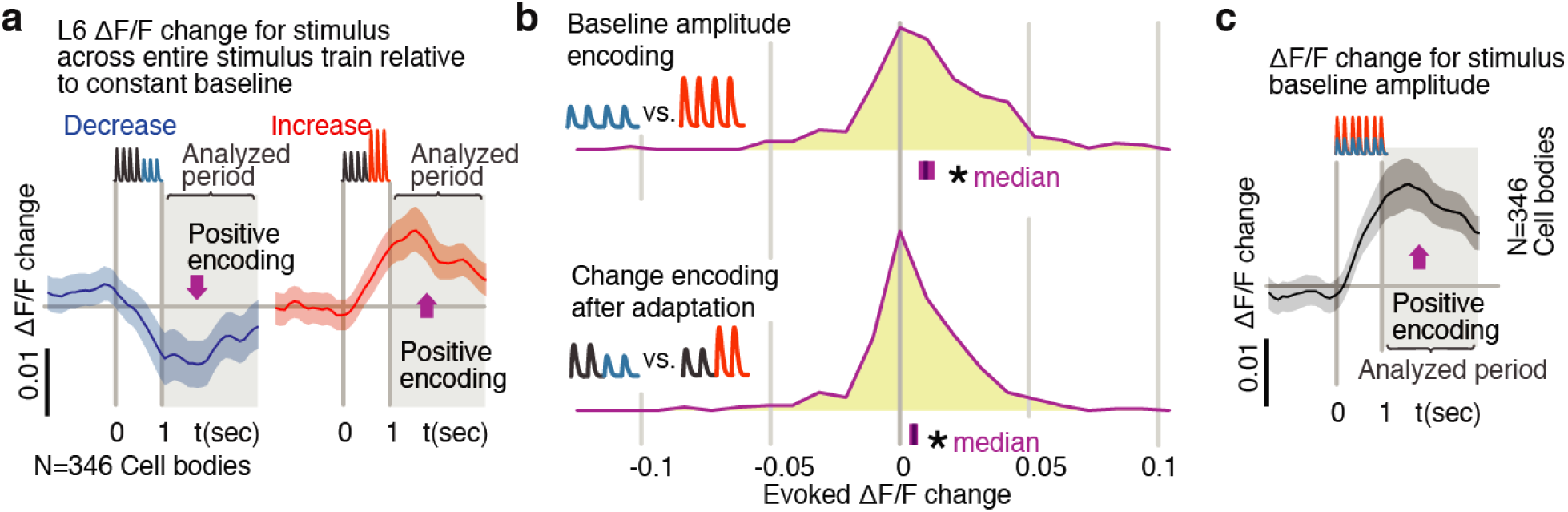
Layer 6 pyramidal cells encoded stimulus amplitude with positive firing rate differences. **(a)** Activity in L6 neurons positively encoded stimulus amplitude, as reflected by larger mean signals among responsive neurons when the response to larger amplitude baseline stimuli was subtracted from the response to smaller baseline stimuli (N = 346 stimulus responsive neurons out of 2685 imaged). **(b)** This positive encoding was also observed when stimulus amplitude variation was provided by deviants later in the stimulus train. *Blue/red* indicate response to increases versus decreases in deviant amplitude relative to baseline. Because calcium imaging data were acquired at ~5 Hz, no attempt was made to quantify change onsets and only the post-stimulus window was analyzed. Bars show the 95% CIs (*purple*) and the median value. **(c)** Population data and summary statistics for L6 CT amplitude and change encoding.

Extracellular recordings in L6 do not yield the sample sizes required to characterize their sensory responses (Fig. 2b) and can not distinguish between corticothalamic (CT) pyramidal cells hypothesized to contribute to change detection, and corticocortical (CC) cells that are less specifically tuned and lack long-range input from higher cortical areas^24^. We therefore employed 2-photon calcium imaging of genetically identified L6 CT cells in GN 220-NTSR1 Cre mice^31^ expressing GCaMP6s^41^ (Fig.2c, N = 3408 cell bodies imaged; N = 2685 tested with amplitude deviants in 6 mice). To study the effect of weak optogenetic drive on the population activity of L6, we combined blue light optogenetic stimulation with 2-photon imaging (Fig.3e, N=1183 cells in 4 mice). Initial attempts to image L6 somata in mice with transgenic GCaMP6s expression, obtained by crossing the NTSR1 Cre and GCaMP6s reporter lines had low signal-to-noise, due to the dense fluorophore expression in the more superficial processes of these cells^42^. To obtain suitable image quality (Fig. 3a-c), sparse labeling of L6 CT neurons was achieved by viral transduction (see Methods). Using this approach, we were able to image throughout the upper ~100 μm of L6 (Fig. 3b-e), and signal-to-noise ratios exceeded 150% ΔF/F (Fig. 3d, Extended Data Figure 7).

**Figure 7.**
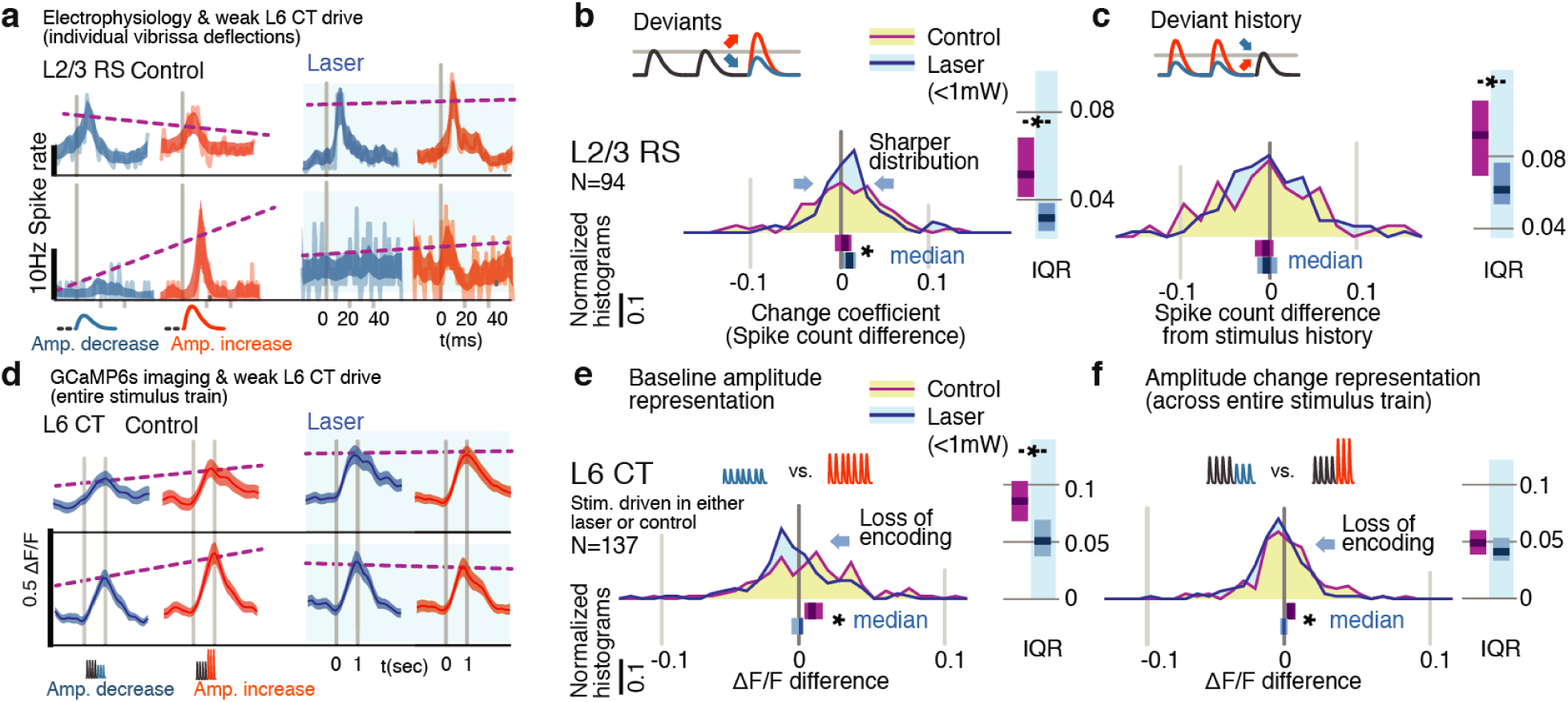
Weak L6 drive disrupts both stimulus encoding in L6 and the emergence of deviant encoding in L2/3. **(a)** L2/3 deviant encoding, of either positive or negative sign, was lost with L6 optogenetic drive. **(b)** Across the L2/3 population, weak L6 CT drive caused a loss in the heterogeneity of L2/3 encoding, reflected as a sharpening of the distribution and bias to positive change coefficients, paralleling L4 encoding. Bar graphs show the 95% CI and the median value for the distribution, and for its interquartile range (IQR), via bootstrap. **(c)** Encoding of stimulus history in L2/3 was similarly disrupted. **(d)** Examples of L6 CT sensory responses (via Ca^2+^ imaging, see Figs.3,6) with and without optogenetic stimulation. In these examples, weak L6 CT optogenetic drive disrupted the representation of stimulus amplitude. **(e)** Population distributions show the loss of baseline amplitude encoding, and **(f)** amplitude encoding for changes within the stimulus trains, in L6 CT.

As in the electrophysiological data, stimulus-evoked L6 activity was sparse (Extended Data Figure 8), and overall L6 calcium signals (relative firing rates) remained unchanged during weak optogenetic drive (Fig. 3f, P=0.10, signed rank). However, weak optogenetic manipulation changed L6 receptive fields: Individual L6 cells showed suppression or facilitation, including both emergence of newly stimulus driven cells and the complete removal of sensory responses (Fig. 3g,h). As such, while the net spiking output of L6 did not change, the specific ensemble of sensory responsive cells changed substantially with this manipulation. As such, this level of optogenetic drive provides a direct approach for perturbing L6 sensory ensembles without detectably altering L6 firing rates.

### Small variations in stimulus strength were not reliably encoded in layers 4 and 2/3 of SI

To understand how shuffling the active ensemble of L6 CT cells selectively affects the ability of mice to detect small sudden stimulus deviations (Fig.1), we next asked how such vibrissa stimuli are encoded throughout the cortical column. We delivered trains of vibrissa deflections (7 deflections, 10 Hz) to the B and C row vibrissae of awake head-fixed mice. To approximate the stimulus deviations in gap-crossing where mice determine distances to objects via small differences in the amplitude and velocity of vibrissa contacts^30,32^, we selected a narrow range of randomized baseline stimulus amplitudes (~25mm/second ±15%), and inserted deviants in stimulus amplitude, ±5-15% of baseline (baseline ~=25mm/sec, N=72 sessions). This stimulus design thus has two parameters: baseline amplitude and deviant amplitude, with deviants varying as an increase or decrease.

We analyzed the firing probability per vibrissa deflection in our chronic extracellular recordings, as a function of baseline amplitude, and deviant amplitude. Variations in stimulus amplitude (without deviants present) were not encoded in the firing rates of layer 4 (L4) or L2/3 RS cells (*L4* N=37 RS; spike count difference between larger and smaller stimuli, signed rank P=0.221, 95% confidence interval (CI): [−0.22, 0, 0.002], *L2/3* N=94, P=0.201, CI:[-0.009,-0.003,0]). Small variations in overall stimulus strength were therefore not reliably encoded in SI.

### Amplitude deviants were not typically reflected by generalized rate increases or decreases

We next asked if deviations in stimulus amplitude following repeated baseline deflections at a fixed amplitude (analogous to a small change in object distance in gap-crossing) were reflected in firing rates. The most commonly reported effect of stimulus deviants is an increased neocortical response^7,10^ regardless of deviant identity. To test for such encoding, we grouped all deviants (increases and decreases) and examined their overall effect on firing rates. While individual neurons displayed sensitivity to deviation (Figure 4f), neither L2/3 nor L4 neurons systematically increased their overall firing rates for deviants (Fig. 4g), showing that our stimulus design avoided pre-cortical stimulus-specific adaptation^7^.

### Layer 4 neurons encoded stimulus deviations with positive change coefficients

We next asked whether stimulus deviants in the middle of an ongoing stimulus (increase vs. decrease) impacted encoding in these layers. Figure 5a shows examples of the types of responses found. Receptive field transformations included significantly greater firing in response to deviants of increasing amplitude (as in examples i and iv), and the opposite, increased responses to deviants that decreased in amplitude (examples ii and iii). To quantify these effects, we calculated the spike count difference between responses to deviants with increased or decreased amplitude, per cell, per deflection. We term this difference a ‘change coefficient,’ with greater firing to amplitude increases referred to as a ‘positive’ change coefficient, and vice versa.

L4 neurons consistently showed positive change coefficients, firing more spikes for increases and fewer for decreases. This encoding is evident at the population level as a rightward shift in the distribution of change coefficients (Fig. 5b, P=0.005, CI:[0.012, 0.016, 0.035]; See Extended Data Figure 11 for generalized linear model ‘GLM’ analysis). These spike count differences for deviations were larger than for equivalent variations in baseline amplitude with no deviant (P=0.016, signed rank). We observed no significant deviant encoding in L5 (N=92: P>0.05). While these effects were small when considered for single neurons and single, small amplitude vibrissal deflections (<0.1 spikes/deflection), they are substantial when considered across populations of neurons, and in the context of vibrissal deflections during natural exploration and gap-crossing. This increased sensitivity of amplitude encoding in L4 after adaptation to a baseline is consistent with the enhanced discriminability of stimulus features generally observed with adaptation^43–46^.

### Layers 2/3 neurons explicitly encoded stimulus amplitude deviations

We next examined change encoding in L2/3 neurons (N=94 vibrissa-driven neurons out of 363 recorded). As in L4, a subset of L2/3 neurons showed positive change encoding, but negative change coefficients were also observed, i.e. neurons with increased firing in response to stimulus decreases (Figure 5a, examples ii and iii). At the population level, such heterogeneity would be reflected in a broadening of the change coefficient distribution relative to non-deviant stimuli. To test whether this heterogeneity was significant at the population level, we computed a surrogate distribution from shuffled baseline stimuli (Fig. 5b, grey). The observed distribution was significantly broader than the surrogate (interquartile range /IQR; Fig. 5c; P=0.006, Shannon entropy: P=0.029, via bootstrapping, see Methods). Ideal observer decoding showed the same difference between positive deviant encoding in L4, and heterogeneous encoding in L2/3 (Extended Data Figure 12). The rate differences for deviants in L2/3 for small stimulus amplitude deviations (± 15%) corresponds to a median difference of spike counts of ≥ ~0.03 spikes per deflection per neuron between the deviant categories. For a 700 ms stimulus at 10 Hz, assuming 300 responsive neurons in the aligned somatotopic representation, this change in rates corresponds to a difference of ~70 spikes per vibrissa, or 250-1000 spikes extrapolated across a typical bout of whisking in sensory decision making such as gap-crossing.

These heterogeneous responses observed in L2/3 could represent tuning for specific patterns of deviations relative to baseline stimuli. However, the higher firing rates we observed for smaller amplitude stimuli are also consistent with individual cells tuned to static stimulus amplitude ranges^47^. To disambiguate these possibilities and test whether L2/3 encoded a true history dependent deviant signal, we analyzed trials where deviant stimulus amplitudes were matched, but deviants were preceded by higher or lower amplitude baseline stimuli (55 sessions, Extended Data Figures 13). Neurons with specific amplitude tuning should show a narrowing of the population distribution around zero change coefficient (as the deviant amplitude itself was constant), whereas encoding of stimulus changes should be reflected in a broadening of the distribution. With fixed deviant amplitude, L2/3 RS continued to represent change from stimulus history with a broadened distribution (Fig. 5d; IQR P=0.004, entropy P=0.014, N=87), showing that L2/3 RS cells encode history dependent heterogeneous change signals.

In sum, L4 neurons encode deviant amplitude, reflected in a positive correlation between their rates and the direction of amplitude change. In contrast, L2/3 neurons showed a variety of responses to deviants, with different receptive fields for specific patterns of baseline and deviant (Fig.5). In contrast to prior reports of increased neocortical excitability with stimulus tuning deviations^7^, neither population showed significantly increased overall rates for small amplitude deviants (Fig.4).

### Layer 6 neurons encoded stimulus amplitude, and weak drive removed this encoding

The low sampling rates of sensory responsive L6 neurons with extracellular recordings did not allow the temporally specific analysis of their tuning for deviance performed on L4 and L2/3. However, the 2-photon imaging data did allow us to ask whether L6 neurons encoded stimulus amplitude in general, and how this encoding might be impacted by the weak optogenetic drive that removed the behavioral benefit of a small stimulus deviant (Fig.1).

To this end, we analyzed integrated activity of L6 CT cells by quantifying responses in a 0-2 second window post vibrissa-stimulus offset. Consistent with electrophysiological data, L6 activity was sparse, as only ~13% of L6 neurons were stimulus driven (469/3408 cells, Extended Data Fig 8). When stimuli without deviants were presented, L6 CT cells encoded baseline amplitude, with significantly higher integrated calcium signals for larger stimuli (P<0.001 signed rank, N=346, Fig. 6b). To test whether this encoding of amplitude persisted after stimulus adaptation, and therefore could contribute to change representation, we presented stimuli with amplitude deviations after 400 ms. As with baseline amplitude variations, L6 CT also encoded these amplitude differences (P<0.001, Fig. 6b). The timescale of these data, collected at 5 Hz sampling rate, did not allow disambiguation between explicit encoding of the current deviant, or a delayed encoding that reacts to stimulus changes over timescales >100 ms. Nevertheless, these data show that L6 CT encoded stimulus amplitude as a variation in relative firing rate, which could serve as a baseline for a change-detection computation.

### Weak drive of L6 CT neurons reduced their stimulus amplitude encoding

Given that weak drive of L6 CT cells did not affect baseline or sensory driven RS firing rates across layers (Fig. 2b), but changed the ensemble of stimulus active L6 CT cells (Fig. 3g,h) and selectively suppressed change detection (Fig. 1f), we next asked how this manipulation affected change encoding of amplitude in L6 CT themselves. In L6 CT, weak optogenetic drive removed their encoding of stimulus amplitude (Fig. 7b,c, P<0.05, ranksum, evoked ΔF/F change between stimuli, control vs. laser). This effect could be explained by a ‘shuffling’ of stimulus driven cells, making otherwise non-driven, and poorly-tuned cells, vibrissa-responsive and vice-versa. Consistent with this hypothesis, cells that were selected to be sensory responsive in the control condition showed decreased average responses in the laser condition (P<0.05, controlled for regression to mean). However, encoding was affected even in L6 CT that were stimulus driven in the control condition and remained so in the laser condition (P=0.044 rank sum across cells, P<0.0001 across trials, Extended Data Figure 9), and the same effect was observed in a cross-validated analysis that classified cells as stimulus driven and analyzed them in separate trials (Extended Data Figure 10). In sum, both the identity of the sensory driven ensemble and amplitude encoding by L6 CT was disrupted by weak optogenetic drive, despite no change in the net activity in these neurons.

### Weak L6 drive removed deviance encoding in L2/3

We next examined whether weak optogenetic drive of L6 impacted stimulus representation in other layers. We found that change representation in L2/3 was disrupted (Fig. 7e, P=0.010 entropy reduction, P**=**0.020 IQR reduction, P=0.008 paired left tailed IQR, N=94). During optogenetic drive, L2/3 neurons came to represent current stimulus amplitudes with positive change coefficients (CI: [0.002, 0.009, 0.016], P=0.003 signed rank) and reduced their history-dependence (Fig.7f, P=0.014 entropy reduction, P=0.004 IQR reduction, P=0.043 paired left tailed IQR). Extended Data Figure 11 shows this same effect quantified via GLM. Shuffling the active L6 ensemble therefore removed the change specific RFs in L2/3, causing them to instead encode current stimuli. These results suggested that the stimulus encoding in L6 is necessary for deviance-specific coding to arise in L2/3.

Recent studies have concluded that optogenetic drive of L6 impacts sensory responses in visual neocortex through an intracortical pathway, and not by modulation of thalamic relay neurons^26,38^. We tested whether weak optogenetic stimulation affected lemniscal neurons using chronic recordings from ventral posterior medial nucleus in awake animals (Methods as in Figs.2,4,5,7). Of 240 well-isolated thalamic single units (N=3 mice), 25 cells were phasically responsive at short latencies to vibrissal stimulation. These neurons showed weak rate increases with L6 optogenetic drive, in contrast to the suppression observed in prior studies using stronger L6 CT drive^35,38^. However, no significant encoding of deviants was observed, nor was modulation of encoding of deviants by L6 drive observed (Extended Data Figure 14). While mechanisms that are not captured by these recordings (e.g., changes in thalamic synchrony) could contribute to the loss of heterogeneous encoding in L2/3 with weak L6 drive, rate changes in thalamic responses were not evident. Further, as described above, layers that receive direct lemniscal thalamic input (L4, L5 and L6) did not show explicit deviance encoding other than encoding of current stimulus amplitude. These two forms of evidence, and prior studies finding a direct intracortical pathway as the primary route of influence by L6 CT^38^, supports a local neocortical transformation in the emergence of L2/3 deviant responses and in their modulation under L6 optogenetic drive.

In sum, the weak optogenetic drive of L6 CT employed here did not drive changes in overall RS firing rates (measured extracellularly, Fig. 2b, and via Ca2+ imaging, Fig. 3f, Extended Data Figures 6,11), in contrast to the suppression observed when using strong optical stimuli in our own data (Fig.1,2) and in prior studies^26^. However, this weak optogenetic stimulus regime did substantially alter sensory encoding by L6 CT. These neurons lost their amplitude encoding, and many individual neurons showed changes in their sensory responsiveness, shifting the specific ensemble of L6 CT activated. Further, L2/3 neurons lost their emergent heterogeneous encoding of deviants, showing instead the positive change encoding observed in L4 under normal conditions (Figure 7).

## Discussion

We observed layer-specific encoding of deviants, and robust behavioral sensitivity to small stimulus deviations. Small stimulus amplitude deviants were correlated positively with firing rates in L4 and L6, with deviant amplitude increases driving higher firing rates. In contrast, heterogeneous encoding, and encoding of stimulus history, emerged in L2/3 neurons. Weak optogenetic drive changed the sensory-responsive L6 ensemble, driving and suppressing subsets of cells, and reduced their information content about the stimulus. This manipulation also removed the encoding of stimulus deviants in L2/3, instead leading to encoding of current stimuli. Detection of small stimulus deviations in the gap-crossing task was also lost with the weak L6 optogenetic drive, without altering basic task performance. These results indicate that stimulus encoding by sparse ensembles in L6 contributes to the neocortical circuit that processes sensory deviation, but is not required for basic sensory function.

While both receptive fields and behavior were altered by the weak optogenetic drive employed here, we found that this manipulation had no significant effect on RS firing rates in L6 or other layers. The manipulation also had no effect on free gap-crossing (Fig. 1). In contrast, using stronger optogenetic drive of these same neurons led to suppressive gain modulation in neocortical response amplitudes^26^ and disrupted baseline sensory sensitivity. The specific impact of the weak L6 manipulation on deviant encoding but not sensory gain (Figs. 2,3), and on deviant-driven sensory behaviors but not basic performance (Fig. 1), indicates that this intervention isolated a network mechanism or computation that is selectively involved in stimulus change processing, but not in processing of repeating, or predictable stimuli. The failure of weak L6 drive to impact baseline behavior is in contrast with several findings showing that relatively subtle optogenetic manipulation in SI, for example induced via similarly weak drive of PV+ interneurons^48–50^ or L4 stellate neurons^51,52^, or direct stimulation of single neurons in other layers^53^, can affect baseline sensory detection behavior.

Studies of sensory deviation typically manipulate stimulus features such as tone pitch^5,54,55^, for which there are pre-cortical tuned populations, and observe higher neocortical firing rates for deviants, likely reflecting recruitment of new pools of neurons tuned for these features. Here, we specifically sought to minimize such pre-cortical stimulus-specific adaptation. In natural perception, relevant stimulus changes could lack feature changes for which there are such populations, e.g. decreases in the amplitude of a stimulus^32^, or higher-order features relayed from other cortical regions. Further, this stimulus design avoids biasing encoding by increases in neocortical drive across all deviants. This lack of pre-cortical stimulus-specific adaptation is evident in the lack of overall increase in firing rates for deviants (Fig. 4g).

L6 CT cells affect cortical activity via the recruitment of local^56^ and trans-laminar^38^ FS mediated inhibition, and through corticothalamic effects^22,35^. The mechanisms by which activity of specific ensembles of active L6 cells influences state or stimulus encoding in superficial layers are still largely unknown, and could also be mediated through a variety of intermediate cell types, layers, and brain areas that were not recorded in the present set of experiments. While the present results directly support a specific role for L6 in the behavioral benefit of deviants, and in the representation of deviation across neocortical layers, further studies are required to determine the circuit, synaptic and cellular mechanisms by which L6 affects neural encoding and behavior.

The encoding of small stimulus changes in L2/3 RS, specifically the history dependence of these responses, is analogous to the emergence of complex temporospatial receptive fields^20,57^ and mismatch encoding^19^ in upper layers of visual cortex. The diversity in L2/3 responses, where both positive and negative change coefficients were observed, could reflect neuron types defined by biophysical characteristics^58^ or projection targets, such as targeting higher somatosensory or motor cortices^59,60^, or may emerge from differential afferent connectivity.

The sparse stimulus encoding observed in L6 CT (Fig. 2b, Fig. 6, Extended Data Figure. 8) could represent either an explicit per-deflection deviant encoding, as in L4 (Fig. 5), or a delayed stimulus or expectation encoding that emerges over timescales greater than 100 ms. The sparsity of L6 CT activity, and their targeting by long-range corticocortical afferents^24^, suggests that they might be gated by inputs from other higher order neocortical areas, as previously proposed^25^. Measurements of rapid responses to individual stimuli with either faster imaging or more comprehensive electrophysiological methods, and more specific network level manipulations than employed here, will be required to disambiguate these hypotheses.

Even though weak L6 CT drive did not alter stimulus-driven firing rates, this manipulation changed the ensemble of sensory driven neurons. This finding suggests that a sparse ensemble of active neurons in L6, with specific connectivity, is required for deviance encoding. There are two types of mechanisms by which this manipulation could lead to the observed disruption of change encoding in L2/3. The specific population of L6 cells active during optogenetic drive was different from the one active in control conditions. This ‘shuffling’ alone could lead to a disruption of stimulus representation in L2/3 because the new set of active neurons would be decoded differently by recipient layers. Additionally, we found that the population of L6 CT cells that is active during optogenetic drive carries less stimulus information than the population that is active in the control condition (Fig. 7, Extended Data Figures 9, 10), suggesting that any decoding of stimulus information from L6 would be impaired under the optogenetic drive condition.

In this study, we employed two kinds of deviants. Variations from strictly repeating baseline stimuli (Fig.4–7, and Extended Data. Figure. 4) and ethologically relevant deviations in the position of a platform in the middle of sampling by freely behaving animals (Fig.1). In the latter case, the stimulus statistics (mainly vibrissa identity, angles, and curvature upon touch) change continuously as the mice approach or retreat from the target platform^30^ (Extended Data. Figure. 2). In both cases, predictive models were at some level formed within the system, driving the change in response patterns and behavior. The similar findings across these paradigms suggests that the mechanism underlying the observed effects could be involved in more general predictive models.

In sum, we found that stimulus encoding by specific ensembles of L6 cells is required for change encoding in L2/3 and for change detection behavior, but not basic detection performance. L6 cells could therefore be one node of the larger laminar cortical circuit for processing of higher order stimulus features, stimulus context, or expectations reliant on top-down signaling, in agreement with L6 CT targeting by long range cortico-cortical inputs^23,24^. (xxx cite other velez fort margrie)

## Online Methods

### Animal subjects

NTSR1-Cre mice (strain B6.FVB(Cg)-Tg(Ntsr1-cre)GN220Gsat/Mmcd, stock number 030648-UCD)^31^ of either sex were used. For some experiments, NTSR1-Cre mice were crossed with a floxed ChR2 reporter line (strain B6;129S-*Gt(ROSA)26Sortm32(CAG-COP4*H134R/EYFP)Hze*/J, stock number 012569). For head-fixed behaviour, one NTSR1-Cre mouse using viral injection, and 3 reporter line crosses (NTSR1/ChR2 +/+) were used. For gap-crossing, 6 NTSR1 mice (2 ChR2 viral injection, 4 reporter line crosses) were used. For electrophysiology, 5 mice of either sex, 4 NTSR1-Cre mice using viral injections and 1 reporter line cross was used. For 2-photon imaging, 6 NTSR-1 Cre mice with viral injections were used.

### Viral injection

For virus mediated ChR2 expression, we targeted the caudal region of the barrel field (1.5 mm posterior to bregma and 3.5 mm lateral to the midline). Injections were performed through a burr-hole with a glass micropipette (pulled and beveled, tip diameter of 20-35 μm) attached to a stereotaxic-mountable syringe pump (QSI Stoelting). 300 nl of virus (AAV DIO ChR2-mCherry; ~2 × 10^12 viral molecules per ml) was injected at 0.05 μl/min at ~800μm below the dura. All experiments requiring viral transfection were performed >4 weeks after injection. For 2 photon imaging, ~300 nl of AAV2/1-hSyn-Flex-GCaMP6s (HHMI/Janelia Farm, GENIE Project; ~1 × 10^13 viral molecules per ml)^41^, or in a subset of mice a 1:1 mixture of floxed GCaMP6 and floxed ChR2, all produced by the U. Penn Vector Core, was injected at ~750μm targeting the posterior c-row barrels, identified by vascular landmarks and confirmed using intrinsic imaging to restrict expression to NTSR1+ neurons. Mice were tested for aberrant expression outside of L6 CT cells either by histology (for behaviour and electrophysiology), or by collecting z-stacks (for 2-photon imaging). Mice with fluorescent non-L6 cells were excluded from the study.

### Surgical procedures

Mice were 8–14 weeks old at the time of surgery. Animals were individually housed and maintained on a 12-h reversed cycle. All procedures and animal care protocols conformed to guidelines established by the National Institutes of Health, and approved by Brown University’s Institutional Animal Care and Use Committee. Mice were anesthetized with isofluorane (2% induction, 0.75–1.25% maintenance in 1 l/min oxygen) and secured in a stereotaxic apparatus. The scalp was shaved, wiped with hair-removal cream and cleaned with iodine solution and alcohol. After intraperitoneal (IP) injection of Buprenorphine (0.1 mg/kg) and dexamethasone (4 mg/kg) and local application of lidocaine, the skull was exposed. For some mice, AAV was injected as described. The skull was cleaned with ethanol, and a base of adhesive luting cement (C&B Metabond) was applied. A 0.5 mm diameter area of the skull over left primary somatosensory cortex was thinned. A stub of fiber-optic cable (0.22 NA, inner diameter of 200 μm, 1.25 mm OD metal ferrule) was glued into place at the side of the craniotomy using transparent luting cement. Custom head posts (www.github.com/open-ephys/headposts_etc) were affixed with luting cement, the incision was closed with VetBond (3M), and mice were removed from isoflurane. Mice were given 3–10 d to recover before the start of water restriction. For electrophysiology, we implanted flexDrives^36^ with 16 stereotrodes (N=2, 17 sessions), or tetrodes (N=3, 58 sessions) made from 12.5 μm polyimide-coated nichrome wire (Kanthal), twisted, heated and gold-plated to 200–400 kΩ impedance. Lateral electrode spacing was 250 μm. Two stainless-steel screws were implanted anterior to bregma to serve as ground. For some mice we injected AAV as described. A craniotomy was drilled over left SI (~1.5 mm posterior, 3.5 mm lateral, ~2.5 mm diameter). A fiberoptic stub was added as for behavioral testing, and a large durotomy was opened. A layer of bacteriostatic surgical lubricant was added, and the drive was lowered at an angle of ~15° and fixed in place using dental cement. After recovery (>3 d), mice were habituated to the setup and electrodes were lowered into the brain (~2hrs between individual electrodes) while noting when each electrode penetrated the brain. Mice were water restricted as described. During the experimental life time of mice, electrodes were advanced to target neocortical layers and maintain recording quality^36^. For 2-photon imaging, titanium headposts were used, the skull around SI was thinned and flattened. A 3 mm craniotomy was made, virus was injected, and a cranial window^61,62^ ‘plug’ was made by stacking two 3 mm coverslips (Deckgläser, #0 thickness (~0.1 mm); Warner; CS-3R) under a 5mm coverslip (Warner; CS-5R), using optical adhesive (Norland Optical #71). The plug was inserted into the craniotomy and the edges of the larger glass were sealed with vetbond (3M) and cemented in place (Extended Data Figure 7). Dura was left intact. Animals were given >3d to recover. We performed intrinsic imaging^63^ to localize the barrel field. For acute recordings (Fig. 2a), the same procedure as for 2-photon imaging was used, but the window was omitted. Mice remained on 0.75–1.25% isofluorane for maintenance, the craniotomy was kept covered in warm saline and a laminar silicone probe (Neuronexus A1×16-3mm-50-177) was lowered into SI.

### Head-fixed behavioral training and stimulus design

Training began >10d after postoperative recovery and at least >7d after onset of water restriction (1 ml/d). Mice were secured to the head-post apparatus, and rested on a platform. Initial training procedure was as described before^48^. White noise (~65dB) was used to mask auditory cues. If mice licked up to 800 ms after the onset of the vibrissae stimulus, water was delivered. There was an additional time-out period of 2 sec for false alarms, and a pre-stimulus delay period (1-4 sec) was gradually introduced, during which licking resulted in a reset of the delay timer. Vibrissae were stimulated with a custom stimulator based on piezoelectric wafers (Noliac CMBP09). Stimulations consisted of deflections with a fast onset velocity and a slower ~80 ms return to baseline with a small (~10% of peak amplitude) negative deceleration period to reduce a 2^nd^ deceleration peak and to reduce the impact of piezo hysteresis. Several vibrissae, centered around the C2 vibrissa, were gripped ~5 mm from the mystacial pad, making sure to grab the same set of whiskers across sessions for the same animals. Amplitudes were calibrated using videography. Piezo elements were replaced if ringing exceeded 10% of the peak amplitude, or if the stimulus amplitude deviated by >5%, or if any hysteresis was measured. Water delivery was controlled by a solenoid valve (Lee Co.), calibrated to give an ~8 μl per opening (30–60 ms). Licking was detected via infrared detectors. After reaching criterion, optogenetic stimulation was added on half of trials. Stimulation started at a variable offset of 0.2 – 1.5 seconds preceding tactile stimulation and persisted for the duration of the stimulus. Laser power was ramped up with a gaussian onset profile lasting ~200ms (Fig. 2b). For the detection behavior, vibrissa stimulus amplitudes were drawn uniformly from a range (~=0-30mm/sec), adjusted manually to maintain performance while probing small stimulus amplitudes. In 10% of trials, maximum amplitude stimuli were delivered. Behavioral experiments were controlled using a custom state machine (www.github.com/open-ephys/behavioral_state_machine) written in Matlab via PCI DIO boards (National Instruments). Mice were weighted daily, and animals that did not consume 1ml of water/session or lost weight were supplemented with water in their home cage several hours after the experiment finished.

### Experimental design for electrophysiology and 2-photon imaging

Mice rested on a styrofoam ball supported by an air cushion. Mice were water restricted and monitored as described, and licked a spout to indicate stimulus detection for reward, but no time-outs, or catch trials were used. Sessions were stopped at signs of animal distress and session durations were increased over the first 2-3 weeks of acclimatization, resulting in sessions of ~2000 trials over ~2h. Stimuli were delivered as described, with deviants of relative amplitude of the deflections between ±10% and ±15%. Amplitudes were calibrated to the range that correspond to ~80-100% hit rate in the behavioral detection task. The interval between stimuli was 3-5s. For 2-photon imaging, mice were not water restricted and stimulus amplitudes were sampled from two baseline stimuli and two deviant conditions (increase to 120% or decrease to 80%) in order to increase statistical power for the sparse L6 responses.

### Gap-crossing behavior

Mice (N=6) were implanted with plastic head-posts and fiber stubs, and water restricted as described, vibrissae on the side ipsilateral to the fiber were trimmed. Two mice also had implants for electrophysiology. The gap-crossing apparatus consisted of two facing platforms^29^ (58mm wide) over a custom LED backlight (650nm). Mice were habituated for 2 days prior to the experiments. On day 3, the optical fiber was attached and masking noise (~80 dB) was introduced. After mice crossed the gap in either direction, a water reward (~0.01 - 0.05 ml) was delivered manually, and a new platform position was chosen between 45 and 65mm. In half the trials the laser was on for >1 sec prior till ~2-3 sec after the crossing. In a subset of trials, one platform was retracted by 2mm within 8 ms via a voice coil actuator (^32^ and Fig.1d, Extended Data Figure 1, and Supplementary video) while the mouse was palpating it. Mice were run every other or third day, sessions ended when mice either lost interest in crossing, fell from the platform, or tangled the optical tether. The gap was filmed from above at 315 Hz (Pike 032B, Allied Vision Technologies). In control sessions, the optical cable was attached to a mock ferrule that directed the light to a position rostral of the actual fiber stub implant.

### Gap-crossing analysis

The mouse nose distance to the target platform was tracked using custom scripts in Matlab. Trials were identified as attempted (mouse reached over the gap) or completed crossings. For analysis of sensory disruption using high laser powers, the probability of crossing was computed from all trials with gap distance <6cm. For other analyses, only trials in which the mice crossed within 5 sec were further analyzed. The nose position over time was aligned to the position at which the mouse had committed to a crossing attempt without touching the target yet, extending over the home platform by ~7 mm, corresponding to a position of -20mm in the imaging reference frame. For analysis of the whisking pattern, subsets of vibrissae were tracked using an automated tracker (Extended Data Figure 2, www.github.com/jvoigts/whisker_tracking) and the median angle of all tracked vibrissae was analyzed.

### Optogenetic stimulation

In all experiments, light was delivered through a jacketed fiber-optic cable 200μm in core diameter and 2.5 m long with a numerical aperture of 0.22 (Doric Lenses) connected to a 450nm diode laser (powertechnology.com) using a collimator (Thorlabs PAF-X-15-PC-A). The fiber was connected to the animal’s head via mating metal ferrules in a zirconia sleeve. For head-fixed behaviour, ferrules were shielded with black plastic tape and the head of the mouse was illuminated with a blue masking LED that did not illuminate the stimulator or vibrissae. Light loss in the implanted fiber stub was measured for each implant. The amplitude of the light stimulus was calibrated regularly with an optical power meter (Thorlabs PM100D with SI20C sensor) to up to 1mW at the surface of the skull, resulting in ~0.1-0.5mW in neocortex (measured through the skull and metabond after perfusion). In a subset of sessions, higher laser power (~2-5mW, see Fig. 2, or up to ~10mW for gap-crossing, Fig. 1) was used. The chronic implantation of a optical fiber results in a somewhat decreased power delivery to the brain due to inevitable regrowth of dura under the implantation site. Direct 1:1 comparisons of the light powers of the chronic experiments to the acute experiments where the fiber was placed directly on the brain (Fig.2a) are therefore not possible, and it should be assumed that a somewhat higher light power is required in the chronic case to achieve the same extent of optogentic activation as in the acute case.

### Analysis of electrophysiology data – acquisition and pre-processing

Unless indicated, we used non-parametric Wilcoxon rank sum / Mann-Whitney U-test tests for comparing groups (non-paired), or Wilcoxon signed rank tests for testing medians versus zero or comparing paired measurements. Extracellular voltage traces were band-pass filtered to 1-10000Hz at acquisition (3^rd^ order butterworth filter), and band-pass filtered to 300-9000Hz (zero-phase acausal FIR) for analysis of spiking. Sessions in which vibrissae had slipped out of the stimulator were excluded. Spikes were sorted into single units using Simple Clust (www.github.com/open-ephys/simpleclust). The 90% quantiles of neuron count/session were 17 and 39 over all 75 sessions. We recorded the depth at which electrodes penetrated the brain, marked by emergence of off-diagonal peak-to-peak amplitudes in the MUA activity (presumably from L1 axons) as the 0mm position to estimate the depth of electrodes. We combined this information with the drive screw position and angle of the drive to estimate depth. In deeper layers, we additionally used the depth at which electrodes entered the white matter (loss of cortical activity) as a further reference point. The mapping from depth to layers was approximately (in μm): L2/3, 100–350; L4, 350–450; L5, 450–650; L6, >650 but was adjusted to take electrode angle and curvature of cortex and white matter borders into account. Drive depth estimates were verified at the L3 /L4 boundary of primary somatosensory cortex (SI), via the stimulus evoked LFP signature^37^.

### Analysis of electrophysiology data – classification of sorted units

We classified neurons as regular spiking (RS) and fast-spiking (FS) by spike waveform^38^. Stimulus driven neurons were classified by fitting a generalized linear model^64^ (GLM) to the PSTH. We classified cells as phasically driven if either of two conditions were met: (i) An offset term and 6 bins (basis functions) spanning the first 100ms of the first vibrissa deflection were fit, coefficients for at least 2 bins were significantly nonzero at a P level of 0.03 and any coefficients other than the offset term had a lower standard error bound > 0.002. (ii) A constant term and 6 repeated bins for over first 100ms of the first 3 deflections were fit, these coefficients were shared between deflections capturing cases of weaker but sustained stimulus drive. Additionally, one parameter for each 100ms vibrissa deflection period after the first one was used to avoid false positives due to slower firing rate drifts. Cells were classified as driven if the coefficients for the first 3 deflections satisfied the same conditions as in (i). Classifications was verified manually to choose thresholds but no manual corrections were made.

### Analysis of electrophysiology data – population coding analysis

To plot example PSTHs (Fig.4,5,7), we computed confidence bounds using a state space method^65^. These analyses were used for visualization purposes only. The random position of deviant stimuli resulted in more baseline than deviant deflections, and more baseline stimuli early in the train (stimuli after the deviants were not analyzed). For analyses that are susceptible to biases of unequal N and adaptation effects, such as change coefficients, a histogram matching procedure was used to match the number and position in the stimulus train across baseline and deviant stimuli (Extended Data Figure 13). The effect of stimulus history on firing rates was analyzed using subsets of trials in which the stimulus amplitudes were matched but were preceded by higher or lower amplitude stimuli by matching the stimulus amplitude distributions (Extended Data Figure 13). All statistics of change coefficients were computed as 95% confidence intervals (CI) of the median using a 1,000 or 10,000-fold bootstrap. Spread of distributions of change coefficients was quantified as the difference between the observed and a surrogate distribution (computed from position matched, randomly re-sampled baseline stimuli) via the interquartile range (75^th^ -25^th^ percentile) and Shannon entropy (in bits): H(observed) - H(surrogate); H(h)=-sum_i(P(h_i) * log2 P(h_i)). The null distribution is computed by re-sampling trials within-cell, and is therefore affected by cell-dependent differences in variability in the same way as the true distribution. 95% confidence bounds and significance levels for these statistics were determined via bootstrap analysis. Entropy was quantified via the difference between pairs of binned distributions, so choice of bin size had no significant effect. Where paired samples per cell were available, as in the effect of the optogenetic manipulation, a bootstrap on the median of the absolute value minus the population median was used: median(absolute(coeff_cell-coeff_population)).

### Analysis of electrophysiology data – per-neuron GLM analysis

We quantified encoding in individual neurons with a GLM. We analyzed parameters for spike count as a function of stimulus deviation, mirroring the direct computation of change-coefficients (Fig.5). The features used in the model were stimulus deviation (−1:decreases, +1: increases), baseline amplitude, and spiking history (for 7 precedent deflections). The adaptation profile was modeled with a separate feature per deflection, linked with a quadratic penalty term on the pairwise difference (weight 10). A separate quadratic regularization term with (weight 1) penalized large parameters (other than constant) to avoid over-fitting. The regularizing matrix (q) was: q=1*I+10*D (I: Identity matrix, D: difference operator). Model parameters (w) were estimated from spike counts (Y) via min_w(−log(p(Y|w)) + 0.5*w’*q’w). 95% confidence bounds were obtained using a 100-fold bootstrap. False positive rates were calculated by fitting to surrogate data (as described). Control, laser, and deviant conditions (increases/decreases) were fitted independently.

### 2-Photon Imaging

A two-photon microscope (Bruker/Prairie Technologies) using an 8 kHz resonant galvanometer (CRS) for fast x-axis scanning, and a non-resonant galvanometer (Cambridge 6215) for y-axis increments was used. In some sessions, non-resonant scanning in a smaller imaging window (variable region ~100×80p×) was used. Frames were 512 × 512 pixels (resonant) or smaller (non resonant) and scanned at >5Hz. Objectives (Nikon 25× 1.1 NA or Nikon 16x 0.8 NA) were rotated to the window plane. GCaMP6s was excited by a pre-chirped Ti-Sapphire laser (Spectra Physics; MaiTai) at 980 nm. Power at the sample was 20-60 mW for superficial imaging (<450μm), 60-80 mW for deep imaging (>450μm), when scanning at ~5-10Hz with an approximate pixel dwell time of 1-2μs. Emitted photons were collected through the imaging path to a multi-alkali PMT (Hamamatsu; R3896, digitized with 14-bit resolution). A typical session lasted 2 hrs. We found no activity ‘run-down’, substantial bleaching or cellular damage over the session, consistent with the what other studies using similar laser intensities have reported^66,67^. In about half of implanted animals, we were able to image cell bodies of NTSR1+ layer 6 CT cells (3408 ROIs total) at depths between ~650-800 μm. Good image quality at commonly used excitation laser powers (see above) at these depths was possible likely due to the sparse and relatively localized expression in L6 (approximate diameter of region with cell bodies ~300-400 μm), which results in relatively little fluorophore above the imaging plane, resulting in better signal to noise ratios at such depths than would be possible with denser labeling^42^. If the optical quality of the implanted window was non-optimal, due to dura re-growth, animal age or any surgical imperfections, L6 imaging became impossible. All analysis routines were written in MATLAB. Motion artifacts, small movements in the x-y plane were corrected with rigid-body image alignment^68^ using a DFT based method^69^ or a similar affine deformation to register to templates averaged from 1000 low-motion frames. To manually identify ROIs, we calculated mean and standard deviation projections, and correlation coefficients for the entire image relative to a seed pixel, and areas of continuous or nearby highly correlated pixels were grouped into the ROI.

### Simultaneous 2-Photon Imaging and Optogenetic stimulation

Light was directed at the entire imaging area from a 200μm fiber at a ~40 degree angle. To minimize light artifacts and PMT damage, we used a blocking filter (Semrock OD 6, custom NIR block, notches to block 460-470nm & 560-570nm). Light from blue (470 nm) or yellow (560 nm) LEDs, driven with a high-speed LED driver (cyclops^70^, designed by Jon Newman, www.open-ephys.org/cyclops) was pulsed for 75μs after each 4^th^ or 8^th^ x-scan line. Overall pulse rates were >200Hz, functionally equivalent to constant light^71^. Light levels were adjusted manually to integrated powers of ~0.1mW (for ChR2). X-scan lines following laser pulses were brighter due to the light stimulation and were replaced by interpolated data from preceding and following x-scan lines, whether the LED was on or off. Remaining slight image brightening was corrected off-line (see below).

### 2-Photon data analysis

Unless indicated, Wilcoxon rank sum or signed rank tests, and bootstrapping for testing IQRs were used, as described for electrophysiological data. Fluorescent values F were extracted from ROIs, the baseline fluorescence F0 was computed as the 30^th^ percentile in a 200 sec sliding window and ΔF/F was computed as (F-F0)/F0. Annulus-shaped ROIs were computed to estimate neuropil contamination^41,68,72^ by eroding out 20 pixels in the x-direction from each somatic ROI (this ensures that if there is any specific artifact from the pulsed optogenetic drive, it affects the cell body and neuropil ROIs equally) and excluding other cell bodies from this neuropil ROI^68,72^. For all analyses of firing rates, residual image brightening due to light artifacts was corrected by subtracting an average image brightening profile averaged from all neuropil ROIs over the entire session. All other analyses are computed as differences in evoked fluorescence between stimulus conditions within the same cell and laser condition, and were therefore not affected by the light artifact correction. Stimulus driven ROIs were identified by comparing the 90% quantiles of the Δ*F/F* for each ROI in the pre-stimulus period for all stimulus conditions (−1500 – 0 ms) with the 10% quantile in the stimulus period (500 - 1500ms), and ROIs with non-overlapping quantiles were analyzed further. Cells were classified either in control trials or optogenetic drive trials, or both. Change coefficients were defined as in the spiking data, but owing to the slow timescale of GCAMP6s, we analyzed the difference of the stimulus evoked fluorescence between baseline and deviant stimuli over the entire post-deviant stimulus time (0-2ec after stimulus offset) instead of analyzing individual deflections. Control levels were computed as described before.

## Data Availability

The datasets generated during the current study are available from the corresponding author on reasonable request.

## Code Availability

All custom software used in this study is freely available: Behavioral experiments were controlled using a custom state machine (www.github.com/open-ephys/behavioral_state_machine) written in Matlab via PCI DIO boards (National Instruments). Vibrissae were tracked using an automated tracker (Extended Data Figure 2, www.github.com/jvoigts/whisker_tracking). Spike sorting was performed using a custom manual sorting tool (www.github.com/open-ephys/simpleclust).

## Acknowledgments

Supported by the NIH: R01NS045130 to CIM and F32MH100749 to CAD. We thank HHMI/Janelia Farm Research Campus and their GENIE Program (V. Jayaraman, R. Kerr, D. Kim, L. Looger and K. Svoboda) for making GCaMP6s available, Tim Buschman for providing behavioural state machine code, and Jon Newman for providing the cyclops LED driver. We thank Scott Cruikshank and Shane Crandall for discussions and comments on the study and manuscript, and Tyler C Brown, Joshua H Siegle and Laura D Lewis for their feedback on the manuscript.

## Contributions

JV and CIM designed the experiments, JV performed electrophysiological and behavioural experiments, JV and CAD performed imaging experiments, JV analyzed the data, JV, CAD and CIM wrote the manuscript.

## Bibliography

1. Chater, N., Tenenbaum, J. B. & Yuille, A. Probabilistic models of cognition: Conceptual foundations. Trends Cogn. Sci. 10, 287–291 (2006).

2. Rao, R. P. N. & Ballard, D. H. Predictive coding in the visual cortex: a functional interpretation of some extra-classical receptive-field effects. Nat. Neurosci. 2, 79–87 (1999).

3. Courchesne, E., Hillyard, S. A. & Galambos, R. Stimulus novelty, task relevance and the visual evoked potential in man. Electroencephalogr. Clin. Neurophysiol. 39, 131–143 (1975).

4. Tiitinen, H., May, P., Reinikainen, K. & Näätänen, R. Attentive novelty detection in humans is governed by pre-attentive sensory memory. Nature 372, 90–92 (1994).

5. Ulanovsky, N., Las, L. & Nelken, I. Processing of low-probability sounds by cortical neurons. Nat. Neurosci. 6, 391–398 (2003).

6. Kutas, M. & Hillyard, S. A. Reading senseless sentences: Brain potentials reflect semantic incongruity. Science 207, 203–205 (1980).

7. Movshon, J. A. & Lennie, P. Pattern-selective adaptation in visual cortical neurones. Nature 278, 850–852 (1979).

8. Chung, S., Li, X. & Nelson, S. B. Short-term depression at thalamocortical synapses contributes to rapid adaptation of cortical sensory responses in vivo. Neuron 34, 437–446 (2002).

9. Katz, Y., Heiss, J. E. & Lampl, I. Cross-whisker adaptation of neurons in the rat barrel cortex. J. Neurosci. Off. J. Soc. Neurosci. 26, 13363–13372 (2006).

10. Khatri, V. & Simons, D. J. Angularly nonspecific response suppression in rat barrel cortex. Cereb. Cortex N. Y. N 1991 17, 599–609 (2007).

11. Knierim, J. J. & van Essen, D. C. Neuronal responses to static texture patterns in area V1 of the alert macaque monkey. J. Neurophysiol. 67, 961–980 (1992).

12. Chelazzi, L., Miller, E. K., Duncan, J. & Desimone, R. A neural basis for visual search in inferior temporal cortex. Nature 363, 345–347 (1993).

13. Reynolds, J. H., Pasternak, T. & Desimone, R. Attention increases sensitivity of V4 neurons. Neuron 26, 703–714 (2000).

14. Maunsell, J. H. R. & Treue, S. Feature-based attention in visual cortex. Trends Neurosci. 29, 317–322 (2006).

15. Diamond, M. E., Huang, W. & Ebner, F. F. Laminar comparison of somatosensory cortical plasticity. Science 265, 1885–1888 (1994).

16. Hyvarinen, J., Poranen, A. & Jokinen, Y. Influence of attentive behavior on neuronal responses to vibration in primary somatosensory cortex of the monkey. J. Neurophysiol. 43, 870–882 (1980).

17. Estebanez, L., Bertherat, J., Shulz, D. E., Bourdieu, L. & Léger, J.-F. A radial map of multi-whisker correlation selectivity in the rat barrel cortex. Nat. Commun. 7, 13528 (2016).

18. Hubel, D. H. & Wiesel, T. N. Receptive fields and functional architecture of monkey striate cortex. J. Physiol. 195, 215–243 (1968).

19. Zmarz, P. & Keller, G. B. Mismatch Receptive Fields in Mouse Visual Cortex. Neuron 92, 766–772 (2016).

20. Martin, K. A. & Whitteridge, D. Form, function and intracortical projections of spiny neurones in the striate visual cortex of the cat. J. Physiol. 353, 463–504 (1984).

21. Zhang, Z. W. & Deschênes, M. Projections to layer VI of the posteromedial barrel field in the rat: a reappraisal of the role of corticothalamic pathways. Cereb. Cortex N. Y. N 1991 8, 428–436 (1998).

22. Thomson, A. M. Neocortical Layer 6, A Review. Front. Neuroanat. 4, (2010).

23. Zhang, S. et al. Long-range and local circuits for top-down modulation of visual cortex processing. Science 345, 660–665 (2014).

24. Vélez-Fort, M. et al. The Stimulus Selectivity and Connectivity of Layer Six Principal Cells Reveals Cortical Microcircuits Underlying Visual Processing. Neuron 83, 1431–1443 (2014).

25. Lee, S., Carvell, G. E. & Simons, D. J. Motor modulation of afferent somatosensory circuits. Nat. Neurosci. 11, 1430–1438 (2008).

26. Olsen, S. R., Bortone, D. S., Adesnik, H. & Scanziani, M. Gain control by layer six in cortical circuits of vision. Nature 483, 47–52 (2012).

27. Bolz, J. & Gilbert, C. D. Generation of end-inhibition in the visual cortex via interlaminar connections. Nature 320, 362–365 (1986).

28. Kok, P., Bains, L. J., van Mourik, T., Norris, D. G. & de Lange, F. P. Selective Activation of the Deep Layers of the Human Primary Visual Cortex by Top-Down Feedback. Curr. Biol. 26, 371–376 (2016).

29. Hutson, K. A. & Masterton, R. B. The sensory contribution of a single vibrissa’s cortical barrel. J. Neurophysiol. 56, 1196–1223 (1986).

30. Voigts, J., Sakmann, B. & Celikel, T. Unsupervised whisker tracking in unrestrained behaving animals. J. Neurophysiol. 100, 504–515 (2008).

31. Gong, S. et al. Targeting Cre Recombinase to Specific Neuron Populations with Bacterial Artificial Chromosome Constructs. J. Neurosci. 27, 9817–9823 (2007).

32. Voigts, J., Herman, D. H. & Celikel, T. Tactile object localization by anticipatory whisker motion. J. Neurophysiol. 113, 620–632 (2015).

33. Celikel, T. & Sakmann, B. Sensory integration across space and in time for decision making in the somatosensory system of rodents. Proc. Natl. Acad. Sci. 104, 1395–1400 (2007).

34. Rema, V. & Chaudhary, R. Deficits in behavioral functions of intact barrel cortex following lesions of homotopic contralateral cortex. Front. Syst. Neurosci. 12, (2018).

35. Denman, D. J. & Contreras, D. Complex Effects on In Vivo Visual Responses by Specific Projections from Mouse Cortical Layer 6 to Dorsal Lateral Geniculate Nucleus. J. Neurosci. 35, 9265–9280 (2015).

36. Voigts, J., Siegle, J. H., Pritchett, D. L. & Moore, C. I. The flexDrive: an ultra-light implant for optical control and highly parallel chronic recording of neuronal ensembles in freely moving mice. Front. Syst. Neurosci. 7, (2013).

37. Castro-Alamancos, M. A. & Connors, B. W. Spatiotemporal properties of short-term plasticity sensorimotor thalamocortical pathways of the rat. J. Neurosci. 16, 2767–2779 (1996).

38. Bortone, D. S., Olsen, S. R. & Scanziani, M. Translaminar Inhibitory Cells Recruited by Layer 6 Corticothalamic Neurons Suppress Visual Cortex. Neuron 82, 474–485 (2014).

39. Swadlow, H. A. & Hicks, T. P. Somatosensory cortical efferent neurons of the awake rabbit: latencies to activation via supra--and subthreshold receptive fields. J. Neurophysiol. 75, 1753–1759 (1996).

40. Helmstaedter, M., Staiger, J. F., Sakmann, B. & Feldmeyer, D. Efficient Recruitment of Layer 2/3 Interneurons by Layer 4 Input in Single Columns of Rat Somatosensory Cortex. J. Neurosci. 28, 8273–8284 (2008).

41. Chen, T.-W. et al. Ultrasensitive fluorescent proteins for imaging neuronal activity. Nature 499, 295–300 (2013).

42. Theer, P., Hasan, M. T. & Denk, W. Two-photon imaging to a depth of 1000 μm in living brains by use of a Ti:Al_2O_3 regenerative amplifier. Opt. Lett. 28, 1022 (2003).

43. Békésy, G. von. Sensory inhibition. (Princeton University Press, 1967).

44. Tannan, V., Whitsel, B. L. & Tommerdahl, M. A. Vibrotactile adaptation enhances spatial localization. Brain Res. 1102, 109–116 (2006).

45. Ollerenshaw, D. R., Zheng, H. J. V., Millard, D. C., Wang, Q. & Stanley, G. B. The adaptive trade-off between detection and discrimination in cortical representations and behavior. Neuron 81, 1152–1164 (2014).

46. Abbott, L. F., Varela, J. A., Sen, K. & Nelson, S. B. Synaptic Depression and Cortical Gain Control. Science 275, 221–224 (1997).

47. Garion, L. et al. Texture coarseness responsive neurons and their mapping in layer 2-3 of the rat barrel cortex in vivo. eLife 3, e03405 (2014).

48. Siegle, J. H., Pritchett, D. L. & Moore, C. I. Gamma-range synchronization of fast-spiking interneurons can enhance detection of tactile stimuli. Nat. Neurosci. 17, 1371–1379 (2014).

49. Lee, S.-H. et al. Activation of specific interneurons improves V1 feature selectivity and visual perception. Nature 488, 379–383 (2012).

50. Sachidhanandam, S., Sreenivasan, V., Kyriakatos, A., Kremer, Y. & Petersen, C. C. H. Membrane potential correlates of sensory perception in mouse barrel cortex. Nat. Neurosci. 16, 1671–1677 (2013).

51. O’Connor, D. H. et al. Neural coding during active somatosensation revealed using illusory touch. Nat. Neurosci. 16, 958–965 (2013).

52. Sofroniew, N. J., Vlasov, Y. A., Hires, S. A., Freeman, J. & Svoboda, K. Neural coding in barrel cortex during whisker-guided locomotion. eLife 4,

53. Houweling, A. R. & Brecht, M. Behavioural report of single neuron stimulation in somatosensory cortex. Nature 451, 65–68 (2008).

54. Taaseh, N., Yaron, A. & Nelken, I. Stimulus-Specific Adaptation and Deviance Detection in the Rat Auditory Cortex. PLoS ONE 6, (2011).

55. Ulanovsky, N., Las, L., Farkas, D. & Nelken, I. Multiple Time Scales of Adaptation in Auditory Cortex Neurons. J. Neurosci. 24, 10440–10453 (2004).

56. Zhang, Z. W. & Deschênes, M. Intracortical axonal projections of lamina VI cells of the primary somatosensory cortex in the rat: a single-cell labeling study. J. Neurosci. Off. J. Soc. Neurosci. 17, 6365–6379 (1997).

57. Gilbert, C. D. Laminar differences in receptive field properties of cells in cat primary visual cortex. J. Physiol. 268, 391–421 (1977).

58. Jouhanneau, J.-S. et al. Cortical fosGFP Expression Reveals Broad Receptive Field Excitatory Neurons Targeted by POm. Neuron 84, 1065–1078 (2014).

59. Sato, T. R. & Svoboda, K. The Functional Properties of Barrel Cortex Neurons Projecting to the Primary Motor Cortex. J. Neurosci. 30, 4256–4260 (2010).

60. Chen, J. L., Carta, S., Soldado-Magraner, J., Schneider, B. L. & Helmchen, F. Behaviour-dependent recruitment of long-range projection neurons in somatosensory cortex. Nature 499, 336–340 (2013).

61. Andermann, M. L., Kerlin, A. M., Roumis, D. K., Glickfeld, L. L. & Reid, R. C. Functional specialization of mouse higher visual cortical areas. Neuron 72, 1025–1039 (2011).

62. Goldey, G. J. et al. Removable cranial windows for long-term imaging in awake mice. Nat. Protoc. 9, 2515–2538 (2014).

63. Polley, D. B., Kvasnák, E. & Frostig, R. D. Naturalistic experience transforms sensory maps in the adult cortex of caged animals. Nature 429, 67–71 (2004).

64. Truccolo, W., Eden, U. T., Fellows, M. R., Donoghue, J. P. & Brown, E. N. A point process framework for relating neural spiking activity to spiking history, neural ensemble, and extrinsic covariate effects. J. Neurophysiol. 93, 1074–1089 (2005).

65. Smith, A. C. et al. State-space Algorithms for Estimating Spike Rate Functions. Intell Neurosci. 2010, 3:1–3:14 (2010).

66. O’Connor, D. H., Peron, S. P., Huber, D. & Svoboda, K. Neural Activity in Barrel Cortex Underlying Vibrissa-Based Object Localization in Mice. Neuron 67, 1048–1061 (2010).

67. Huber, D. et al. Multiple dynamic representations in the motor cortex during sensorimotor learning. Nature 484, 473–478 (2012).

68. Bonin, V., Histed, M. H., Yurgenson, S. & Reid, R. C. Local diversity and fine-scale organization of receptive fields in mouse visual cortex. J. Neurosci. Off. J. Soc. Neurosci. 31, 18506–18521 (2011).

69. Guizar-Sicairos, M., Thurman, S. T. & Fienup, J. R. Efficient subpixel image registration algorithms. Opt. Lett. 33, 156–158 (2008).

70. Newman, J. P. et al. Optogenetic feedback control of neural activity. eLife 4, e07192 (2015).

71. Lin, J. Y., Lin, M. Z., Steinbach, P. & Tsien, R. Y. Characterization of Engineered Channelrhodopsin Variants with Improved Properties and Kinetics. Biophys. J. 96, 1803–1814 (2009).

72. Kerlin, A. M., Andermann, M. L., Berezovskii, V. K. & Reid, R. C. Broadly tuned response properties of diverse inhibitory neuron subtypes in mouse visual cortex. Neuron 67, 858–871 (2010).

