## Supplementary Figures for "Layer 6 ensembles can selectively regulate the behavioral impact and layer-specific representation of sensory deviants"

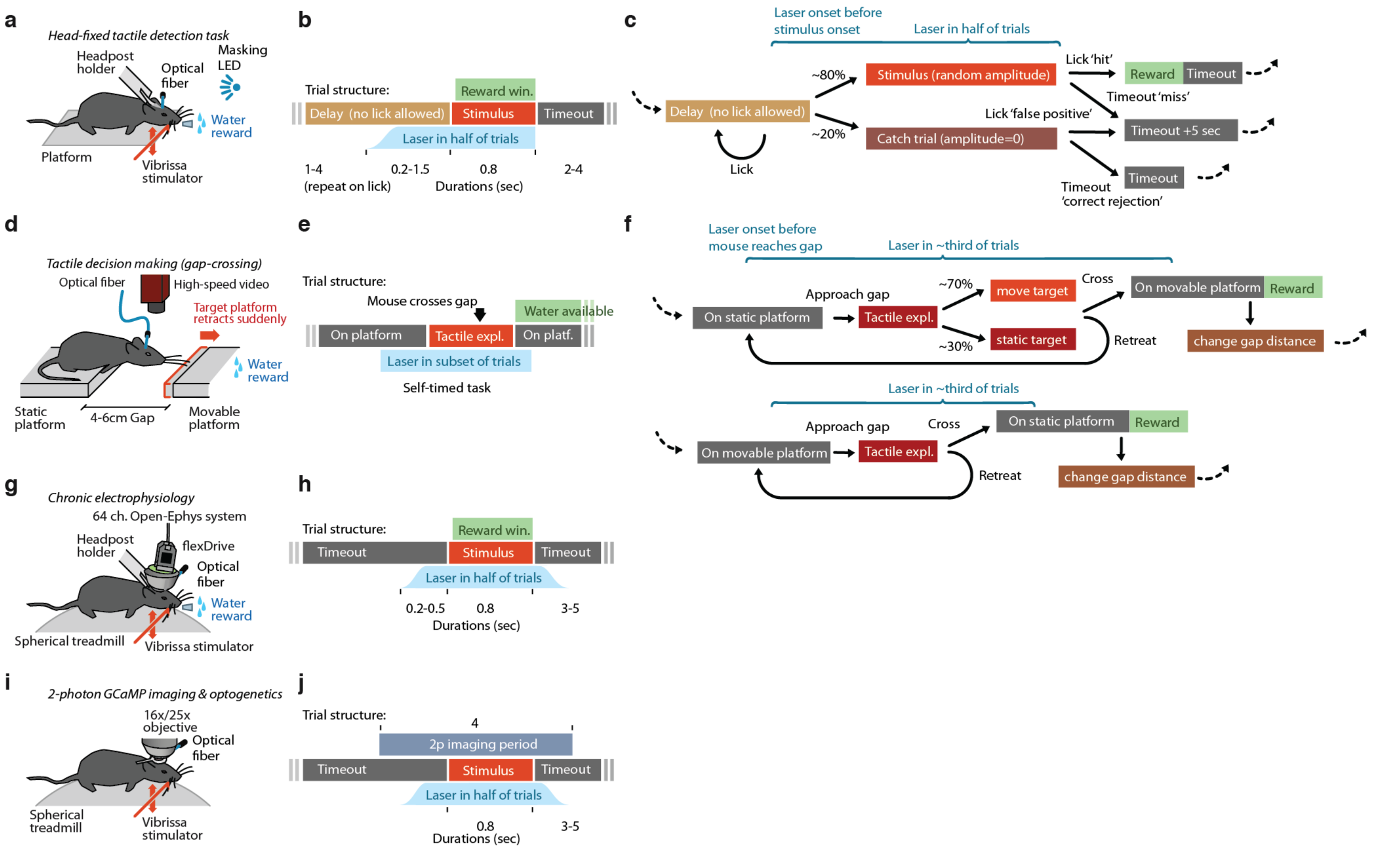


**Supplementary Figure 1 | Overview of trial structure and timing for the experimental paradigms used in this study. a,** Experimental preparation for head-fixed detection behavior (Supplementary figure 4, N=4 mice). Mice were head-posted and rested on a platform. Correct stimulus detection was rewarded with water. **b,** Trial timing for head-fixed behavioral testing. Early licking in a variable timeout period reset the timeout. **c,** State diagram for head-fixed detection task. **d,** Setup for unrestrained gap-crossing behavior (Fig.1, N=6 mice). Mice freely crossed between two platforms for water reward. The platform distances were chosen randomly between 4-6cm, and on a subset of trials the target platform was pulled back by ~2mm mid-exploration. **e,** Timing structure for gap-crossing. All events except for the timing of the laser stimulus and the gap-repositioning were chosen freely by the animals. **f,** State diagram for gap-crossing. **g,** Setup for chronic electrophysiology (Figs. 2,4,5,7, N=5 mice). Correct stimulus detection was rewarded with water. Mice rested or walked/ran on a spherical treadmill to promote comfort and longer recording sessions. **h,** Trial timing for electrophysiology. No timeout periods or catch trials were used. **i,** Setup for 2-photon imaging (Figs. 3,6, N=2 mice) and simultaneous 2-photon imaging and optogenetics (N=4 mice). Mice were not water restricted and rested or walked/ran on a spherical treadmill. **j,** Trial timing for imaging experiments. No reward was used, but stimulus timing was randomized as in other conditions.

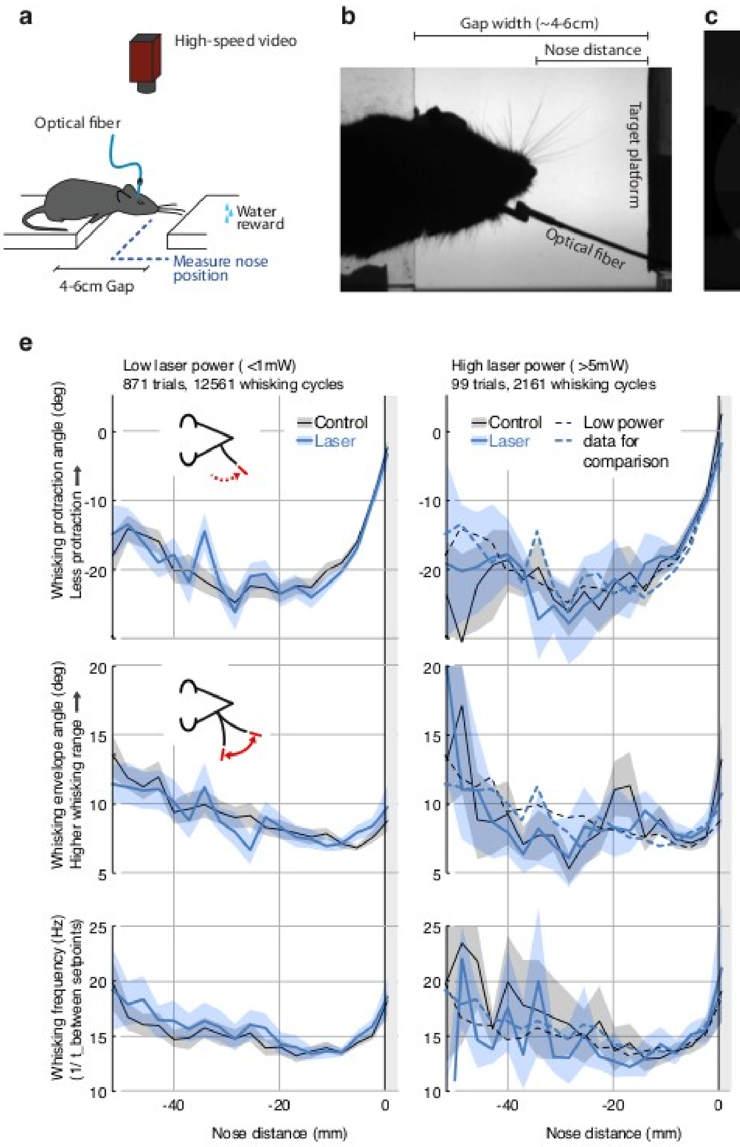


**Supplementary Figure 2 | Whisking pattern kinematics in the gap-crossing task are not impacted by laser stimulation.** Neither the low-power (<1 mW) nor the high-power (>5 mW) optogenetic L6 CT drive caused significant disruption of the sensorimotor whisking pattern in the gap-crossing task. **a,** Overview of setup for gap-crossing[1–4]. **b,** Example raw image from gap-crossing experiment. **c,** Output of the convolutional neural network / vibrissa-identification stage of the vibrissa tracker (github.com/jvoigts/whisker_tracking). **d,** Same as panel c, but individual vibrissa segments are identified with a Hough transform. **e,** Vibrissa kinematics, resolved by nose-target platform distance (N = 4 mice with vibrissa tracking, 970 trials total) for the low-power (<1mW ChR2 drive) and high power (>5mW) conditions. All plots are median and 95% confidence bounds for the median via bootstrapping. The low-power medians (left) are re-plotted as dotted lines in the high power condition (right). Top row: maximum protracted angle per vibrissa protraction cycle. Values closer to 0 correspond to smaller vibrissal protractions. Middle: Whisking pattern envelope amplitude (Hilbert-transform on 8-20 Hz filtered median whisker angle). Bottom: Whisking frequency computed from time between whisking pro/retraction setpoints. Data points above 30 Hz were excluded.

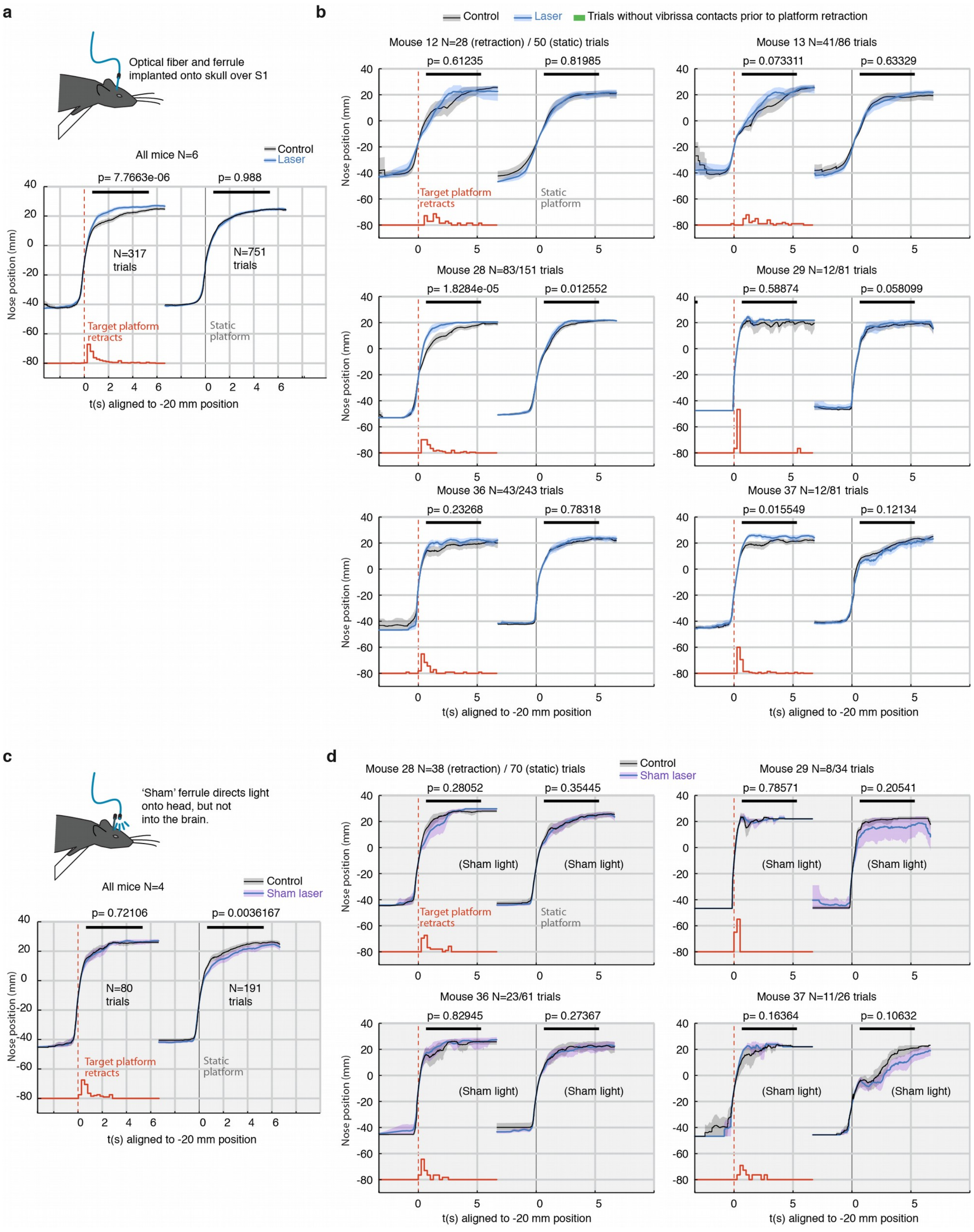


**Supplementary Figure 3 | Per-mouse analysis of freely behaving gap-crossing and sham-laser control experiment. a,** Summary statistics for gap-crossing in the low-power (<1 mW laser) condition, same as Fig. 1. Left: Trials with retracting target (retraction times indicated in red, sum-normalized histogram). Right: return trials (opposite direction) with a static target platform. P-values are from a rank sum test using the mean position for each trial in the region indicated with a black line. Only trials where vibrissae overlap the platform before and after the platform retraction were included because only these trials present sensory change to the mouse. This analysis was performed using fully automated vibrissa tracking and no explicit collision detection other than intersection of vibrissae and target edge was used. **b,** Analysis for the gap-crossing experiment for each individual animal. **c,** Same as panel a, but for sham-laser condition. Mice (N = 4) were tested as usual, but a 'sham' ferrule directed the laser light diffusely over the entire head of the animal instead of coupling it into the implanted ferrule. These control experiments were run with higher laser powers of ~10 mW to ensure that the light was always brighter and more visible to the animals than in the main experiment. **d,** Analysis for the sham-laser experiment for each individual animal. There was no specific effect on stimulus change detection.

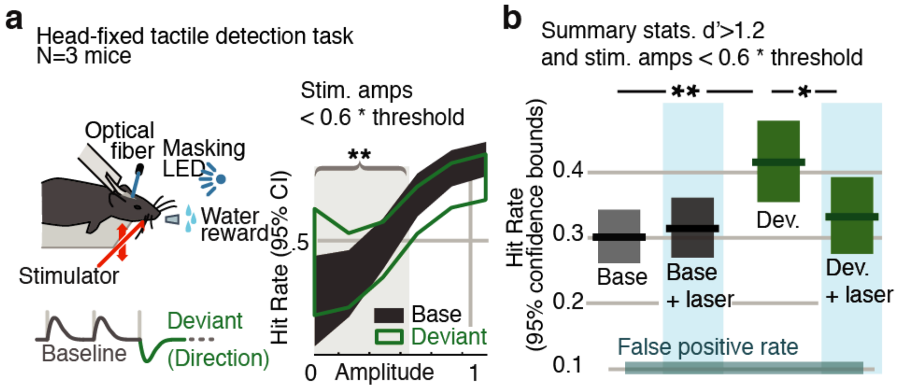


**Supplementary Figure 4 | Weak activation of L6 negates the improved sensitivity driven by inclusion of deviant stimuli.** Stimulus deviations can improve the detection of sensory stimuli[5,6]. Particularly, the detection of less-salient, harder to perceive stimuli are robustly impacted by dynamics in vibrissal SI[7]. We tested the impact of weak L6 drive on this enhanced sensitivity using deviations in the direction of vibrissal motion[8]. This stimulus design differs from the amplitude deviations used throughout the rest of the study, but the more salient direction deviants are well suited to cause increased stimulus salience, and therefore make it possible to observe a behavioural effect in a detection task. **(a)** Mice were trained to indicate stimulus detection for water reward[7,9–11] (go/no-go task). Mice wer enot explicitly trained to report the presence of a deviant. Mice that learned the task (N = 3) were run for 50-100 sessions, and periods of high performance were analyzed (d' statistic > 1.2). In 50% of trials, direction deviants were present at positions 2-4 in the stimulus train. See Supplementary Figure 15 for control condition for auditory or visual cues. As shown in the psychometric curve from one mouse, inclusion of deviants increased detection rates for weak stimuli that were less than 60% of the threshold amplitude (*gray region*). **(b)** Box plots show 95% CIs (via bootstrap) for the detection probabilities for this range of stimuli from 3 mice. Inclusion of a deviant increased the detection rate (*green* bar labeled 'Dev.'). However, when weak L6 CT optogenetic drive was applied, no effect was observed on baseline detection rates, but deviants no longer caused significantly enhanced sensitivity. This change-specific behavioral effect of weak L6 drive was accompanied by a loss of deviant-specific encoding in L2/3 and 4, quantified via electrophysiology, and in L6, via 2-photon imaging (Supplementary Figure 16).

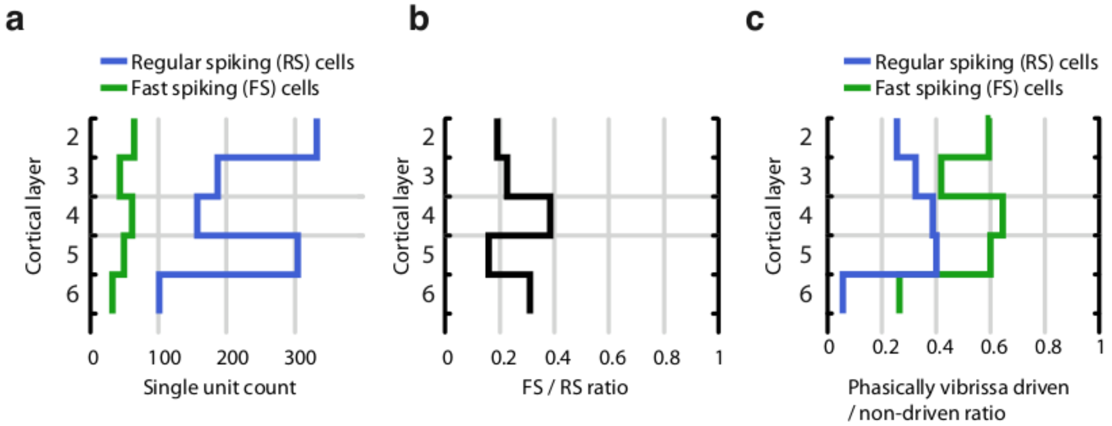


**Supplementary Figure 5 |** Proportion of fast spiking (FS) and regular spiking (RS) neurons across cortical layers. a, Total per-layer count of RS and FS cells. b, Proportion of FS to RS cells across all layers. c, Proportion of significantly vibrissa-stimulus driven (see Methods) RS and FS neurons across layers. The proportion of stimulus-driven cells is biased by our attempts to record from stimulus-driven cells – tetrodes were adjusted when recording conditions had stabilized but no stimulus-driven activity was recorded. Therefore, our data set likely significantly over-represents stimulus driven cells.

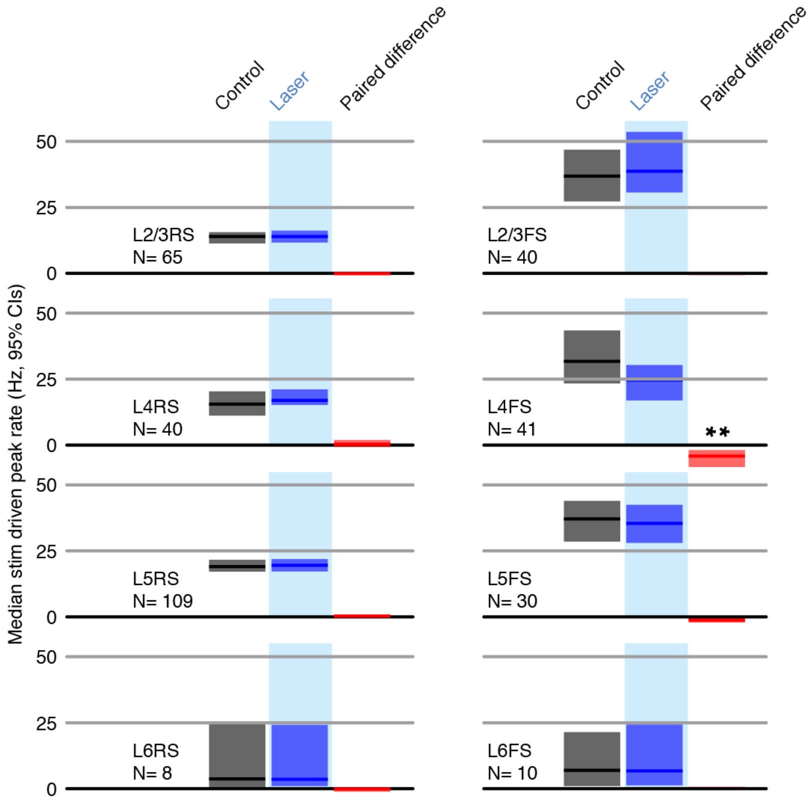


**Supplementary Figure 6 | Peak firing rates (peak responses for deflections 2-7 for each neuron) for control and laser conditions across all stimuli.** All shaded error bars are 95% confidence intervals (CI) of the median.

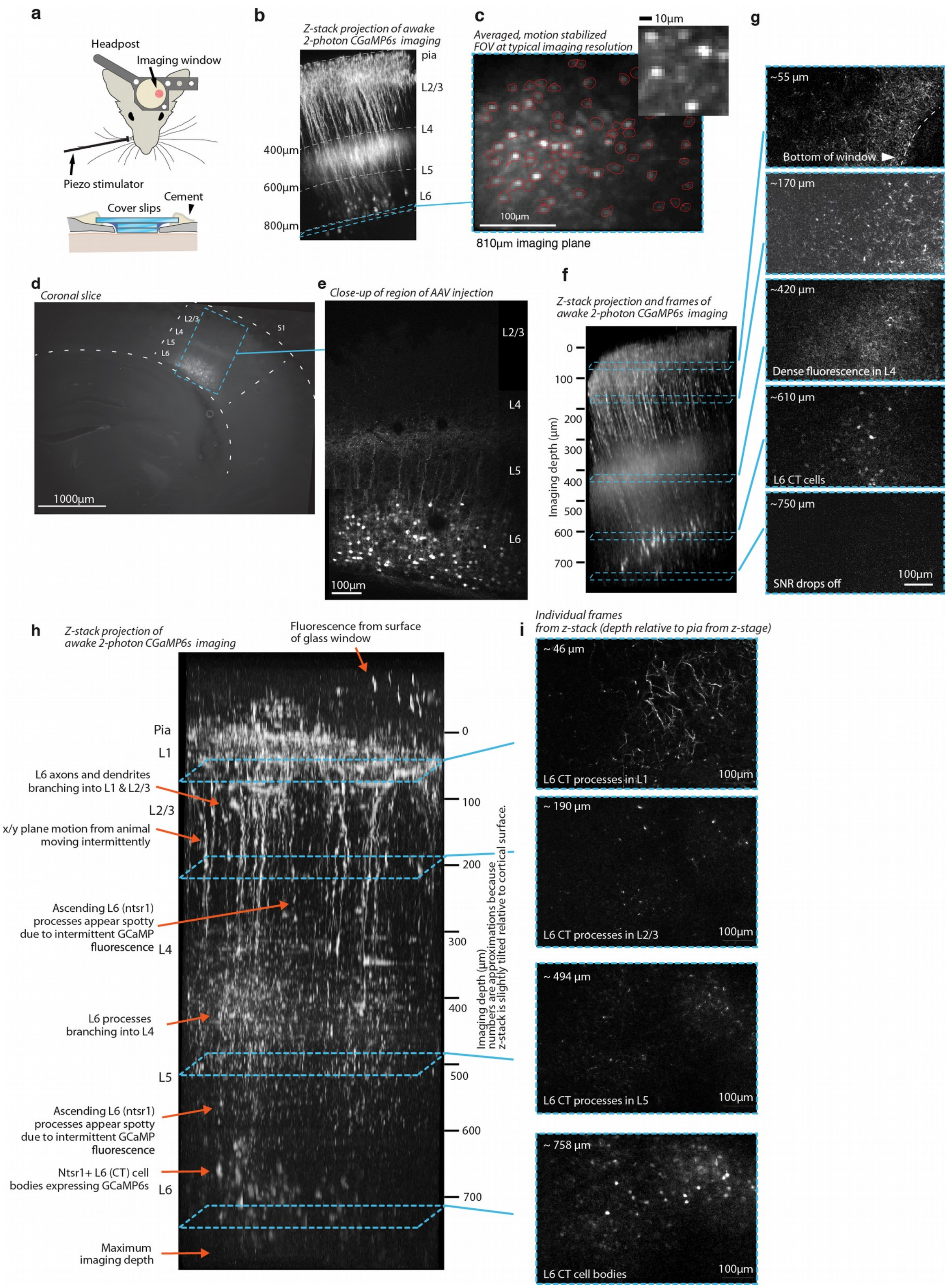


**Supplementary Figure 7 | Histology and 2-photon imaging in L6 CT cells with GCaMP6s in the NTSR1-Cre line. a,** Illustration of the imaging setup using two stacked cover slips under a bigger top cover slip. **b,** Z-stack projection of GCaMP6s image stack acquired in awake animal. **c,** Averaged, motion corrected field of view from typical L6 imaging session. **d,** Coronal section (150µm thickness) imaged at 525nm showing AAV mediated expression of GCaMP6s in NTSR-1 positive L6 CT neurons. **e,** Close-up of same slice (collage of 2 images), acquired with 2-photon scanning. **f,** Z-stack assembled from 170 images at 5µm steps, shown slightly oblique (~10 degrees) to the imaging Z-axis, maximum intensity projection. The stack was acquired in a resting, awake animal. Overall features of the image are identical to panel a and b, but superficial fluorescence appears brighter due to the strong image intensity falloff at greater depths in the 2-photon z-stacks. Beady appearance of processes in superficial layers is due to intermittent calcium driven fluorescence that makes individual dendrites/axons show up as very bright, large spots in individual frames (see panel e, 2nd example frame). Note that the relatively weak and sparse fluorescence above L4 is much more visible here than in the slice. This lack of superficial fluorescence is likely one of the features of the NTSR1 line that make this deep 2-photon imaging possible[12]. **g,** Individual frames from Z-stack in panel f, each image is remapped to equalize contrast and brightness and slightly smoothed to improve visibility (gaussian filter, sigma=0.3px). **h,** Z-stack, same procedure as in c, assembled from 165 images at 5μm steps. **i,** Individual images from the z-stack in panel h.

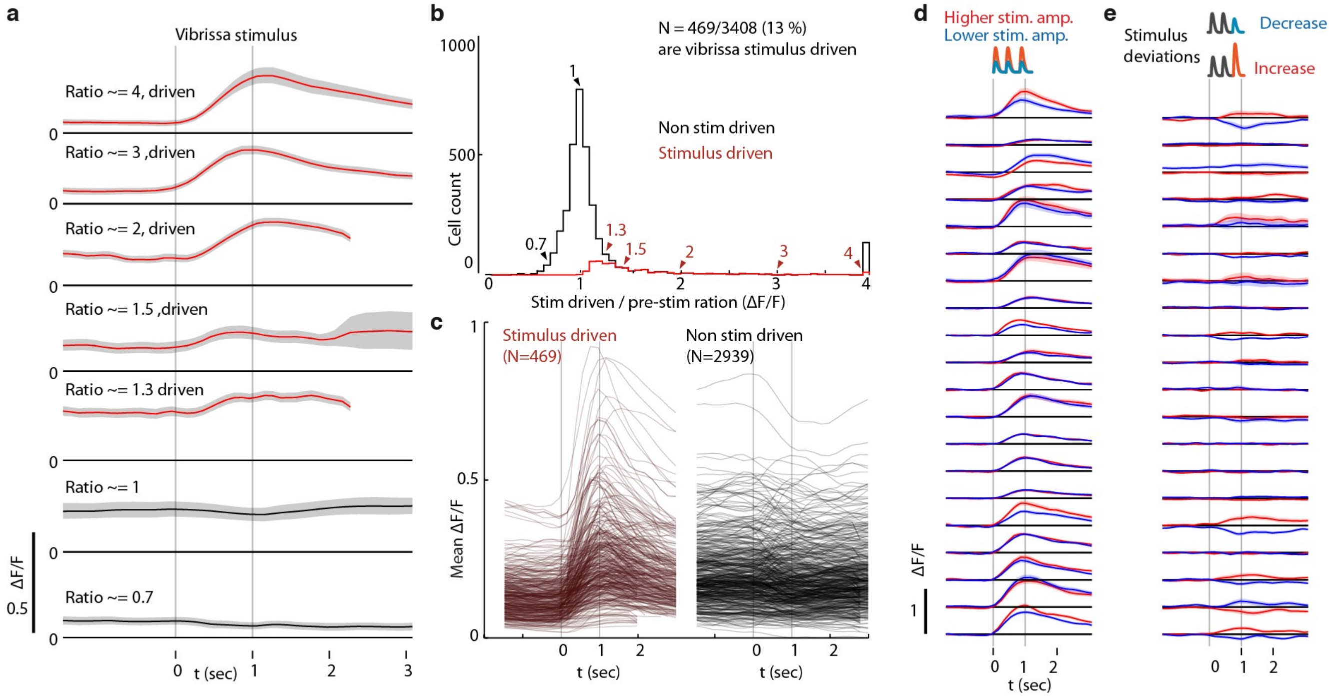


**Supplementary Figure 8 | Classification of L6 CT cells as vibrissa-stimulus driven.**

**a,** Responses (stimulus triggered mean ΔF/F) for 7 example ROIs showing approximate stimulus drive ratios of 0.7, 1, 1.3, 1.5, 2,3, and 4. Stimulus drive amplitude was quantified as the ratio of the ΔF/F in a response window (0-2 seconds after vibrissal stimulus offset) divided by the baseline ΔF/F (-2 to 0 sec). Confidence bounds are 95% for mean via bootstrap. **b,** Histogram of stimulus drive ratios for significantly vibrissa stimulus-driven (red) and non-driven (black) ROIs. ROIs were labeled as significantly stimulus driven if the 25th percentile of the mean ΔF/F in the response window was larger than the 95% percentile of the pre-stimulus period. **c,** Mean responses for all stimulus-driven, and the first 500 non-significant ROIs. **d,** Mean responses of highly stimulus-driven cells for high (red) and low (blue) (~120% or 80% of a mean stimulus amplitude) non-deviant vibrissa stimuli. **e,** Mean difference between constant amplitude vibrissa stimuli and stimuli containing stimulus increase deviants (red) or decrease deviants (blue). Deviants were in the 80%-120% of baseline range.

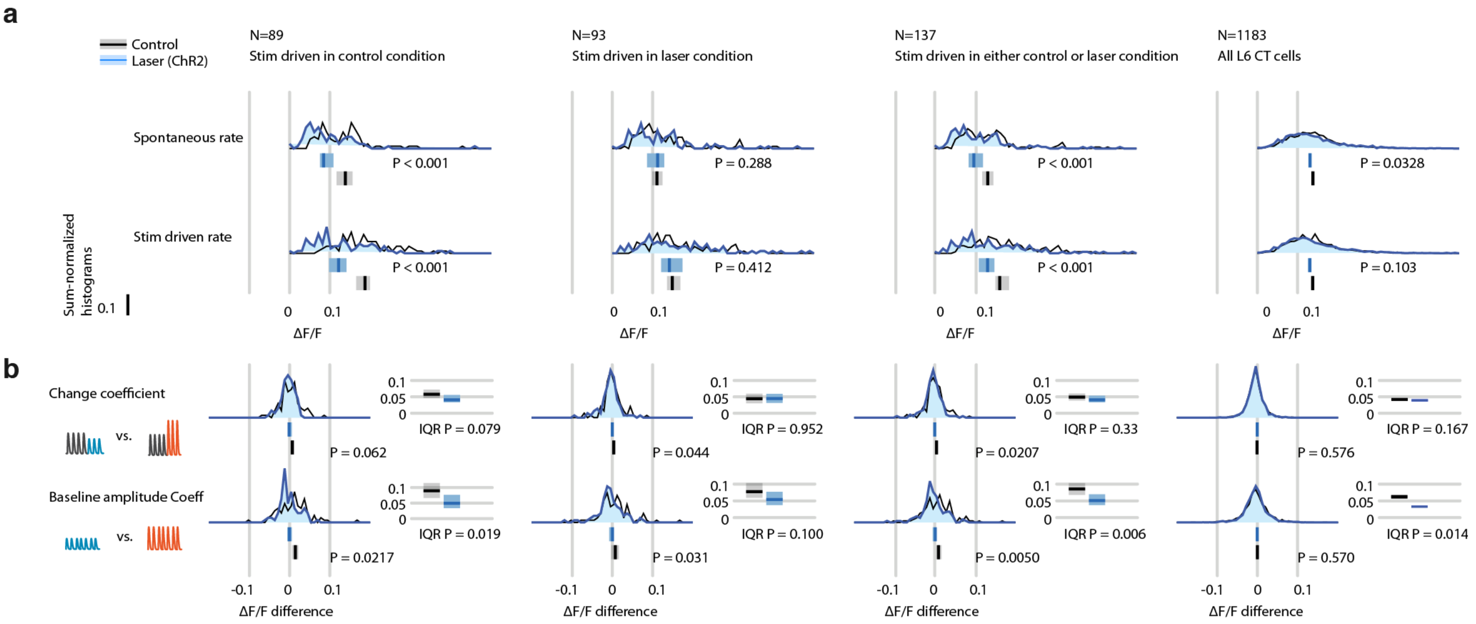


**Supplementary Figure 9 | Weak optogenetic L6 CT drive with ChR2 disrupts encoding of small stimulus amplitude changes and deviants, quantified via raw ΔF/F.** Effect of weak L6 CT ChR2 drive (~0.1-0.2 mW through imaging window, see Fig. 3) on L6 CT cell firing rates during vibrissa stimulation, measured by calcium imaging via GCaMP6s in stimulus driven cells. All vibrissa stimulus responses were quantified in a 0-2 sec window relative to stimulus offset (or equivalent time in catch trials). **a,** Histogram of spontaneous (catch trials) or vibrissa stimulus evoked mean ΔF/F. Bar graphs show 95% CIs of the median per group via bootstrapping (1000 fold). P values are from two-sided Mann-Whitney rank sum tests between the control and laser conditions. Data is analyzed as all stimulus driven cells classified in control condition (left), classified in laser trials (middle left), in either condition (middle right), and for all L6 CT cells (far right). **b,** Histograms of change coefficient, computed as the difference in evoked ΔF/F between constant and increasing amplitude and constant and decreasing amplitude trials, and baseline amplitude encoding (mean difference in ΔF/F between low and high-amplitude constant stimuli). Analysis is split into groups as in panel a. P values are either from Mann-Whitney rank sum tests (unpaired) or Wilcoxon sign rank test (paired by cell). Spread of the distributions is quantified via IQR, CIs are computed via 1000 fold bootstrap, P values via rank sum test.

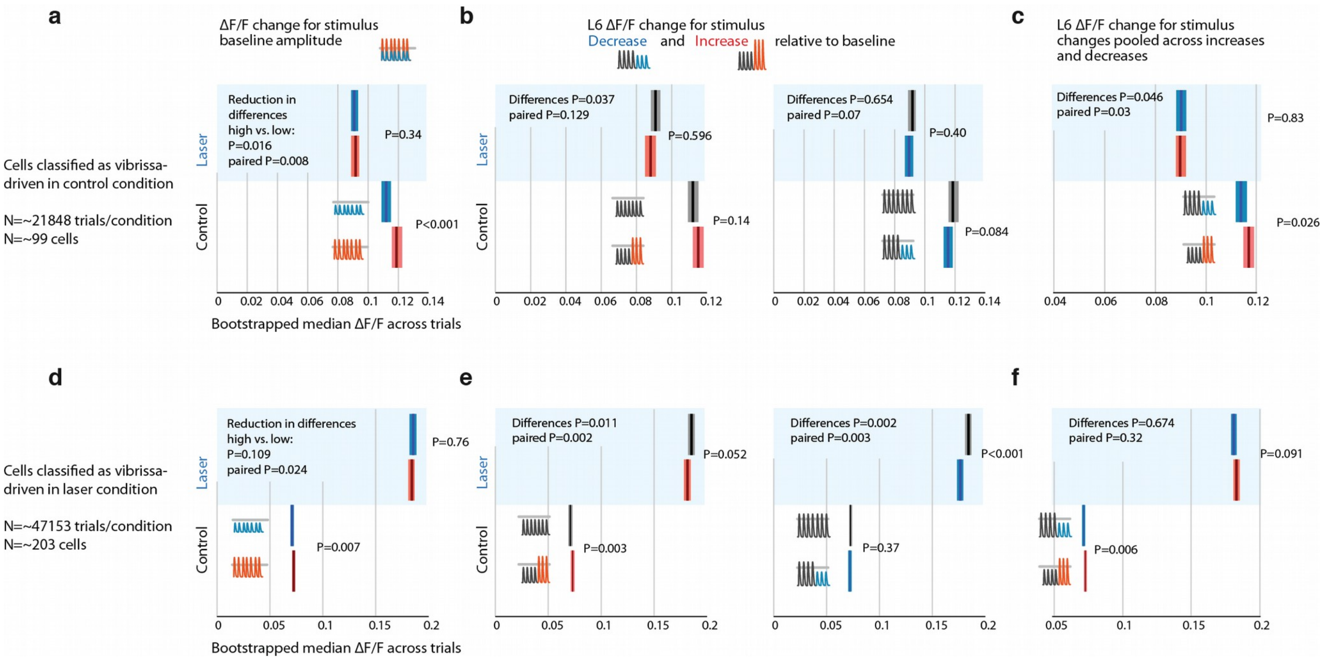


**Supplementary Figure 10 | Weak optogenetic L6 CT drive with ChR2 disrupts encoding of small stimulus amplitude changes and deviants, quantified via leave-one-out cross-validation for stimulus driven vs. non-driven cells.** Effect of weak L6 CT ChR2 drive (~0.1-0.2 mW through imaging window, see Fig. 3) on L6 CT cell firing rates during vibrissa stimulation, measured by calcium imaging via GCaMP6s in stimulus driven cells. All vibrissa stimulus responses were quantified in a 0-2 sec window relative to stimulus offset (or equivalent time in catch trials). **a,** To test whether the encoding of stimulus features in L6 CT cells was affected by weak L6 drive independently of a possible selection bias in choosing cells that are stimulus driven in the control condition and then possibly lose significance in the laser condition purely because of the variance in their responses, we used a leave-one-out (LOO) method to quantify L6 CT cell responses (also used for quantifying the directional tuning on L6 CT cells, see Supplementary Figure 18). Instead of assigning cells as stimulus or not stimulus driven, we classified a cell as stimulus driven per trial, using all but one trial, and then analyzed this trial only if the other trials showed significant stimulus drive. This procedure avoids selecting data on the same dataset as is used for the analysis. In this analysis, positive change coefficients were observed (small stimuli led to smaller, bigger stimuli to bigger ΔF/F). P values for comparisons within control or laser conditions are from rank sum tests. This difference is significantly reduced in the laser condition. P values for the decrease of this difference, in control versus laser trials are computed by bootstrapping across trials and testing the median of the difference (laser vs. control) of differences (large in control-small in control) – (large in laser-small in laser) versus zero for each sample. For paired P values by cells, each bootstrap sample computed differences of the difference of mean responses as before, but within cells, for a bootstrapped sample of cells and then tested the median across cells versus zero. **b,** Comparison of ΔF/F for constant and changing stimuli for increasing (red) and decreasing (blue) trials. **c,** Same data as in panel d but pooled across increasing and decreasing deviant conditions. Deviants were significantly encoded and this encoding was disrupted in the laser condition. Because of the timescale of GCaMP6s, multiple interpretations exist for this encoding. This encoding may be determined by a delayed response to repeated presentation of the deviant amplitude and does not necessarily represent a true deviant encoding. **d,e,f** Same as panels c, d, e but cells were classified as stimulus driven (via LOO procedure) in the laser condition. Baseline stimuli are significantly encoded in these cells, and this encoding is significantly disrupted in the laser condition. Stimulus deviants are encoded in the control condition but the encoding is not reduced as clearly in the laser condition.

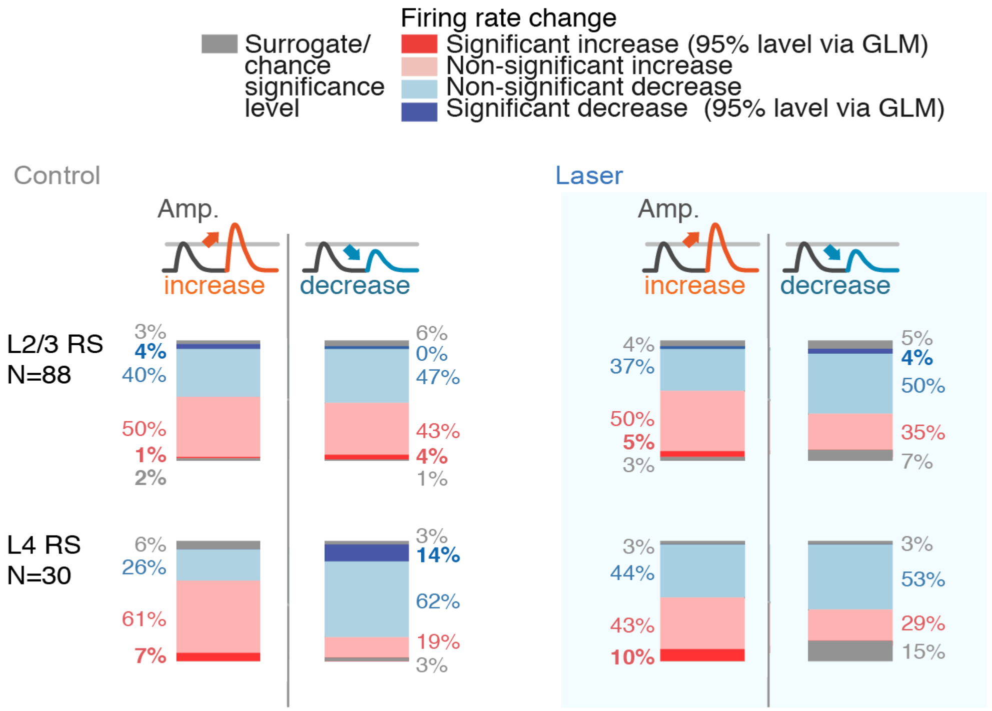


**Supplementary Figure 11 | GLM coefficient statistics across layers and conditions.** Percentages of neurons with significantly increased or decreased firing rates, split by stimulus increases or decreases (each group rounded to nearest integer, as such totals do not necessarily add to 100%). Same analysis was run for control and laser conditions. The shift away from heterogeneous deviant encoding in L2/3 RS in the laser conditions is reflected in the higher proportion of positive coefficients. Significance of individual neurons was assessed using a GLM (see Supplemental Methods), using a 95% significance level at ~500 trials. The (relatively small) numbers of individually significant neurons using this criterion do therefore not reflect the information content of their spike output, but should be interpreted only in their difference across layers, cell types and control versus laser conditions. In L2/3, population-level analysis of deviant encoding shows robust encoding of deviants via the variance of their change coefficients (Figure 2), and via ideal observer analysis (Supplementary Figure 5). As shown in Supplementary Figure 33, relatively small changes in firing rate per cell correspond to robust stimulus encoding across the population of sensory responsive neurons.

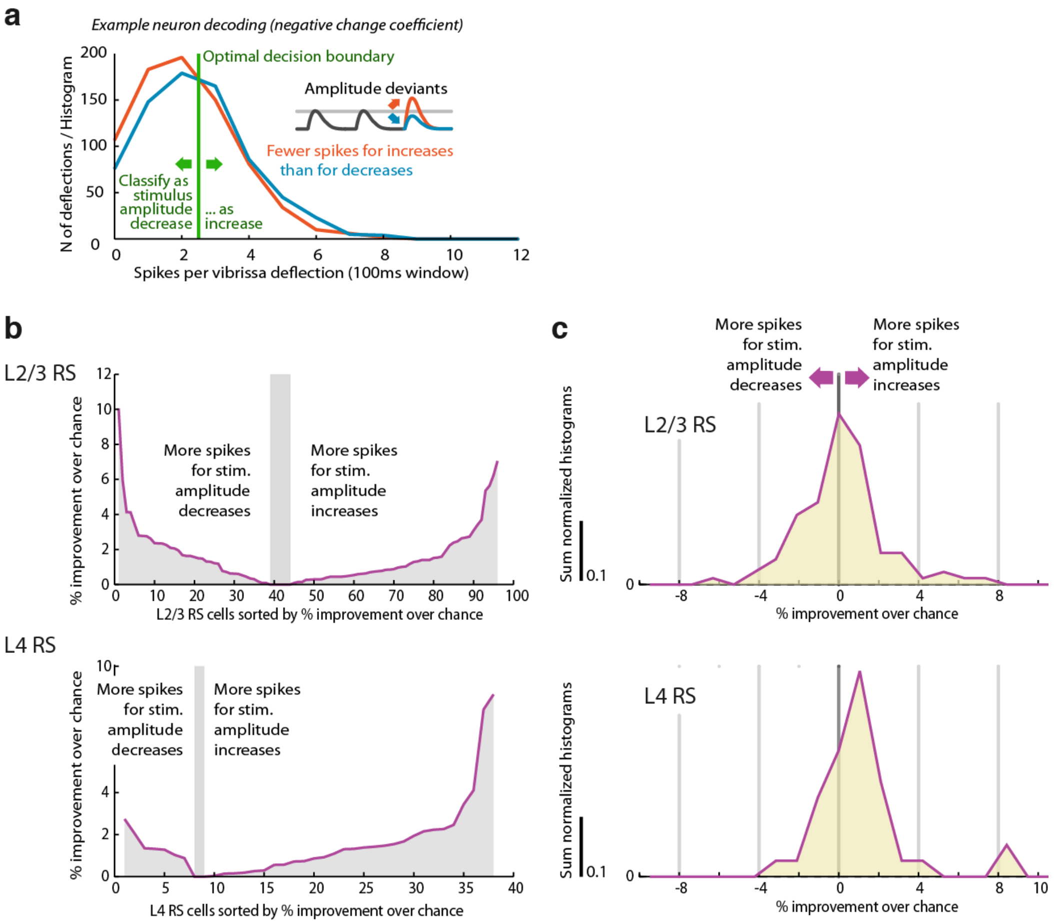


**Supplementary Figure 12 | L2/3 RS cells encode stimulus amplitude deviant identity heterogeneously, in contrast to L4.** L2/3 has an equal proportion of cells with increased or decrease firing probability for stimulus amplitude increases, in contrast to L4, which mirrors the same finding obtained by directly analyzing firing probability for increase/decrease deviants (Fig. 5). **a,** Example of ideal observer decoding for one cell. A optimal spike threshold is chosen as the value that results in the best classification rate (maximally different from 50%). Higher spike counts were always decoded as stimulus increases. For cells that decrease their firing rate for increased stimulus amplitudes or vice versa (i.e., negative change coefficient neurons), the correct rate would therefore be <50%. A cell that fires on every trial with a significantly positive change coefficient could theoretically result in a 100% classification rate, while such a cell with negative change coefficient could yield an (equally informative) 0% hit rate. **b,** Rectified (always positive) correct classification rates relative to chance rate (which can slightly differ from 50% because of unequal proportions of stimulus increase versus decrease trials). In L4, most cells showed positive classification rates / spike count increases for stimulus increases, while in L2/3 the proportions are closer to even. **c,** Histogram of (non rectified) correct rates relative to chance level across cells – the distribution of cells with positive or negative change encoding mirrors that of the change coefficients (Fig. 5).

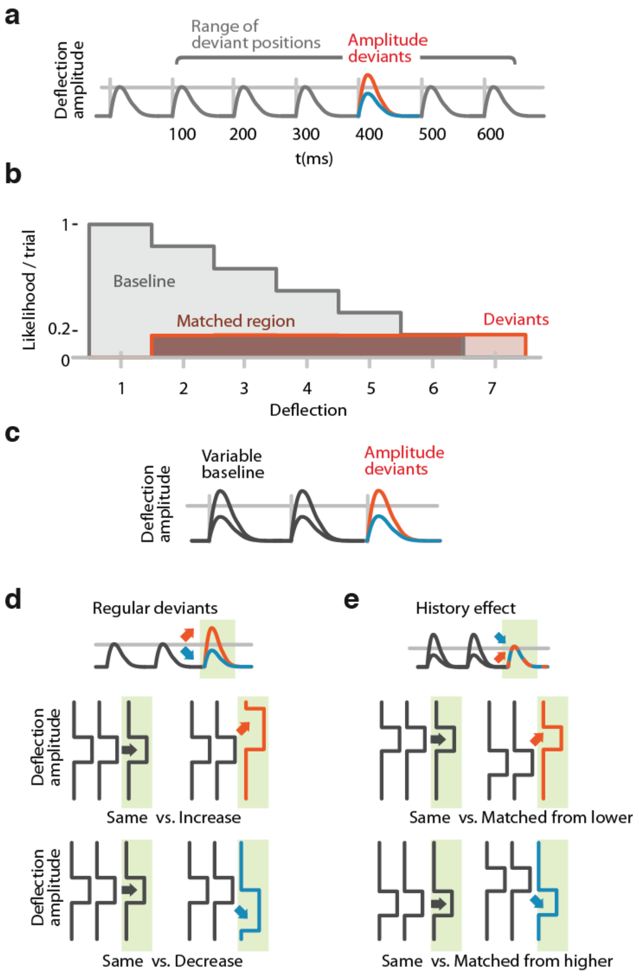


**Supplementary Figure 13 | Method for matching positions in stimulus trains and for analysis of encoding of stimulus history. a,** Possible positions of deviations in the stimulus train at deflections 2-7. **b,** Distributions of positions in trains of baseline (grey) and deviant (red) stimuli. Baseline stimuli are more likely to occur at the beginning of the train (post-deviant stimuli were excluded from analysis). Overlapping region (dark brown) shows the subset of trials that was sampled from to compute surrogate distributions to compare baseline and deviant stimuli while controlling for possible effects of different N and different positions in train. **c,** Stimulus design for sessions in which baseline amplitude was randomized across trials (N=55 sessions). **d,** Selection of trials for analysis of neural encoding of stimulus changes. Green: analyzed stimuli. Baseline stimuli are compared to increased or decreased stimuli. For analyses that are affected by adaptation or different sample sizes, positional matching (see a, b) was used. **e,** Selection of trials for analysis of stimulus history representation. Green: analyzed stimuli. Trials were sub-sampled to obtain conditions in which baseline amplitudes (grey) and deviant amplitudes (red/blue) were matched. The deviant condition was therefore defined by higher or lower preceding stimuli, allowing an analysis of the effects of stimulus history independent of the current stimulus amplitude. For analyses that are affected by adaptation or different sample sizes, positional matching (see a, b) was used in addition to the amplitude matching.

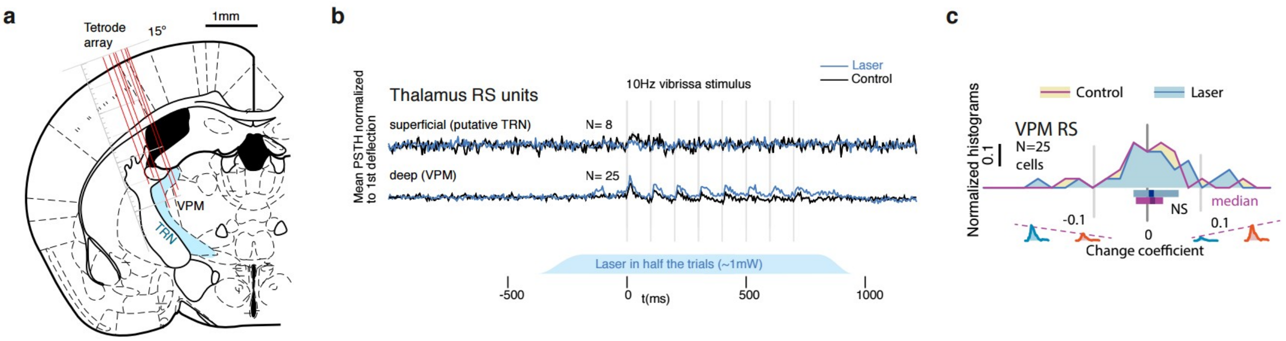


**Supplementary Figure 14 | Thalamic relay cells do not encode stimulus deviants as significantly as cortical L4 RS cells. a,** We recorded a total of 240 putative VPM cells. Recording locations in VPM and neighboring TRN are shown for one animal. **b,** Out of the sample of VPM cells, 25 cells were significantly phasically stimulus driven. We also observed 8 TRN neurons with weak stimulus - onset specific firing. **c,** Same analysis as Fig.5 – Neither in the control condition or under weak L6 drive are stimulus deviants encoded in the recorded VPM cells. TRN cells were not phasically stimulus driven past the first deflection and carried no stimulus information.

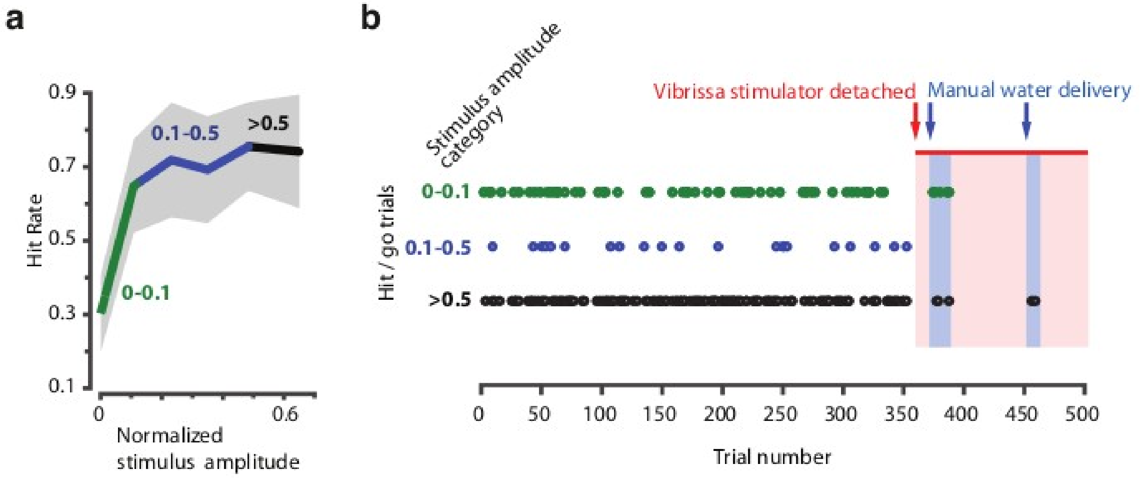


**Supplementary Figure 15 | Head-fixed behavior depends on vibrissa stimulation and is independent of other (auditory or visual) cues.** Mice were trained to detect vibrissa stimulation (in a separate experiment) using the same vibrissa stimulator, masking noise, absence of <650nm light, and behavioral protocol as describes in Supplementary Figures 1 and 4. **a,** Psychometric curve for one session, stimulus amplitudes were split into three categories based on stimulus amplitude (<0.1, 0.1 – 0.5 and >0.5 x maximum amplitude). **b,** Raster plot of successful detections (licks in go-trials) over the session (500 trials). Misses or catch trials are not plotted. Towards the end of the session, the stimulator was opened freeing the vibrissae, but remained in the same position, so that vibrissa contact to the stimulator was still possible and any vibration, visual or auditory cues were the same as in the beginning of the session. In two blocks, reward was given manually to verify that the animal was still attentive and could react to reward delivery (indicated by the click of a solenoid) with licking (indicated as hits in cases where water was given in non-catch/go trials).

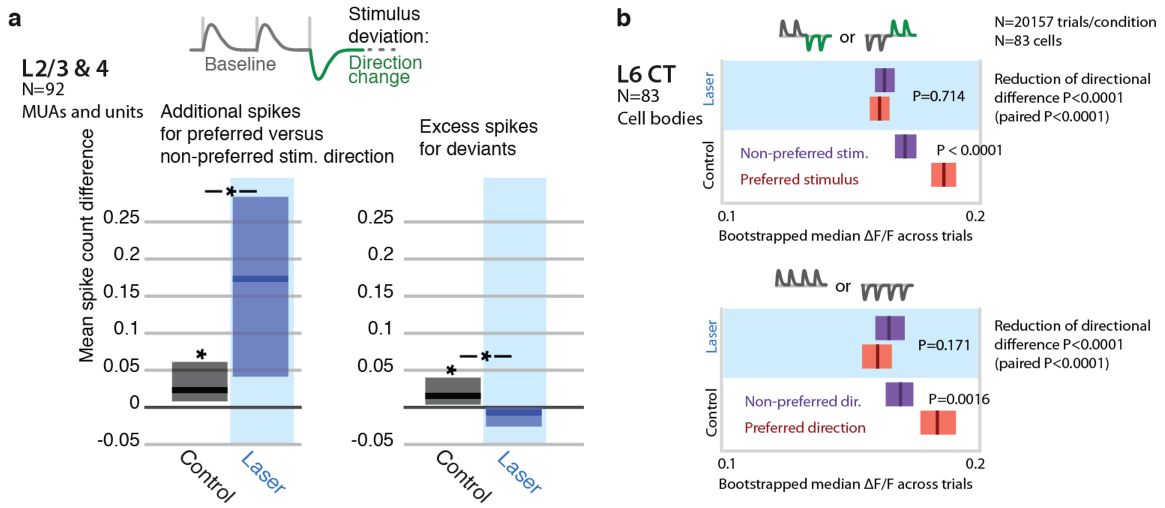


**Supplementary Figure 16 | Encoding of directional deviants, and effect of L6 activation.** The direction changes employed for the head-fixed detection task (Supplementary Figure 4) differ from the amplitude deviants used throughout the rest of the study. We therefore also examined the impact of weak L6 CT drive on the neural representation of direction deviations in separate recordings. In L6 CT (GCaMP6s imaging, N = 83 sensory responsive neurons), encoding of stimulus direction was disrupted by weak optogenetic drive (P<0.05, baseline direction and direction changes). Similarly, L2/3 and L4 neurons (N=92 units and MUA recordings) became more sharply tuned to preferred directions during optogenetic activation (P=0.038 left tailed, change in spike count for preferred over non-preferred), rather than increasing firing rates for deviants (P=0.003 right tailed decrease). These findings parallel the effects observed using amplitude deviants. **a,** Stimulus design for electrophysiological study of directional deviants, same as for head-fixed behavioral stimulus detection task (9 sessions). Baseline direction was randomly chosen (see Methods). Spike rate increase to preferred deviant direction (N=92 units and MUAs) relative to baseline. Preferred direction was determined by the tuning to baseline stimuli. L6 activation increased the relative response to the preferred versus non-preferred direction (rank sum, p=0.038 left tailed increase in spike count increase for preferred). Spike rates were increased for directional deviants (relative to baseline, directions are balanced). L6 drive removes this change encoding (p=0.003 right tailed decrease in additional spikes). **b,** Direction encoding in L6 CT cells assessed using simultaneous 2-photon imaging and optogenetics, (83 cell bodies, GCaMP6s, see main text and methods). Top: L6 CT cells preferentially responded do either trains of upwards, followed by downwards deflections or vice-versa (P<0.0001 rank-sum, 8 deflections, 10Hz, see Methods). For each trial, all other trials of that L6 CT cell body in the same session were analyzed to determine the preferred stimulus and the held out trial was counted either as preferred or non-preferred for the analysis (see Supplementary Figures 9,10 for details**).** Bottom: Same analysis, but whole trains of vibrissa deflections were made up of upwards or downwards deflections. L6 CT significantly encoded direction (P=0.0016). Weak L6 drive disrupted the encoding of deflection direction in L6 CT cells, in both cases (reduction of differences across preferred and non-preferred between control and laser conditions P<0.0001 via non-paired bootstrap across trials in both cases, P<0.0001 paired by cell via bootstrap.

Bibliography for Supplementary figures:


1. Hutson, K. A. & Masterton, R. B. The sensory contribution of a single vibrissa's cortical barrel. *J. Neurophysiol.* **56**, 1196–1223 (1986).
2. Celikel, T. & Sakmann, B. Sensory integration across space and in time for decision making in the somatosensory system of rodents. *Proc. Natl. Acad. Sci.* **104**, 1395–1400 (2007).
3. Voigts, J., Sakmann, B. & Celikel, T. Unsupervised whisker tracking in unrestrained behaving animals. *J. Neurophysiol.* **100**, 504–515 (2008).
4. Voigts, J., Herman, D. H. & Celikel, T. Tactile object localization by anticipatory whisker motion. *J. Neurophysiol.* **113**, 620–632 (2015).
5. Goble, A. K. & Hollins, M. Vibrotactile adaptation enhances amplitude discrimination. *J. Acoust. Soc. Am.* **93**, 418–424 (1993).
6. Musall, S. *et al.* Tactile frequency discrimination is enhanced by circumventing neocortical adaptation. *Nat. Neurosci.* **17**, 1567–1573 (2014).
7. Miyashita, T. & Feldman, D. E. Behavioral Detection of Passive Whisker Stimuli Requires Somatosensory Cortex. *Cereb. Cortex* **23**, 1655–1662 (2013).
8. Khatri, V. & Simons, D. J. Angularly nonspecific response suppression in rat barrel cortex. *Cereb. Cortex N. Y. N 1991* **17**, 599–609 (2007).
9. Stüttgen, M. C. & Schwarz, C. Psychophysical and neurometric detection performance under stimulus uncertainty. *Nat. Neurosci.* **11**, 1091–1099 (2008).
10. Sachidhanandam, S., Sreenivasan, V., Kyriakatos, A., Kremer, Y. & Petersen, C. C. H. Membrane potential correlates of sensory perception in mouse barrel cortex. *Nat. Neurosci.* **16**, 1671–1677 (2013).
11. Siegle, J. H., Pritchett, D. L. & Moore, C. I. Gamma-range synchronization of fast-spiking interneurons can enhance detection of tactile stimuli. *Nat. Neurosci.* **17**, 1371–1379 (2014).
12. Theer, P., Hasan, M. T. & Denk, W. Two-photon imaging to a depth of 1000 μm in living brains by use of a Ti:Al_2O_3 regenerative amplifier. *Opt. Lett.* **28**, 1022 (2003).
